# Scalable multiple whole-genome alignment and locally collinear block construction with SibeliaZ

**DOI:** 10.1101/548123

**Authors:** Ilia Minkin, Paul Medvedev

## Abstract

Multiple whole-genome alignment is a challenging problem in bioinformatics. Despite many successes, current methods are not able to keep up with the growing number, length, and complexity of assembled genomes, especially when computational resources are limited. Approaches based on compacted de Bruijn graphs to identify and extend anchors into locally collinear blocks have potential for scalability, but current methods do not scale to mammalian genomes. We present an algorithm, SibeliaZ-LCB, for identifying collinear blocks in closely related genomes based on analysis of the de Bruijn graph. We further incorporate this into a multiple whole-genome alignment pipeline called SibeliaZ. SibeliaZ shows run-time improvements over other methods while maintaining accuracy. On sixteen recently-assembled strains of mice, SibeliaZ runs in under 16 hours on a single machine, while other tools did not run to completion for eight mice within a week. SibeliaZ makes a significant step towards improving scalability of multiple whole-genome alignment and collinear block reconstruction algorithms on a single machine.

## 1 Introduction

Multiple whole-genome alignment is the problem of identifying all the high-quality multiple local alignments within a collection of assembled genome sequences. It is a fundamental problem in bioinformatics and forms the starting point for most comparative genomics studies, such as rearrangement analysis, phylogeny reconstruction, and the investigation of evolutionary processes. Unfortunately, the presence of high-copy repeats and the sheer size of the input make multiple whole-genome alignment extremely difficult. While current approaches have been successfully applied in many studies, they are not able to keep up with the growing number and size of assembled genomes (Earl *et al.*, 2014). The multiple whole-genome alignment problem is also closely related to the synteny reconstruction problem and to the questions of how to best represent pan-genomes. There are two common strategies to tackle the whole-genome alignment problem (Dewey and Pachter, 2006). The first one is based on finding pairwise local alignments (Altschul *et al.*, 1990, 1997; Schwartz *et al.*, 2003; Harris, 2007; Kent, 2002) and then extending them into multiple local alignments (Blanchette *et al.*, 2004; Dubchak *et al.*, 2009; Angiuoli and Salzberg, 2011; Paten *et al.*, 2011a). While this strategy is known for its high accuracy, a competitive assessment of multiple whole-genome alignment methods (Alignathon, Earl *et al.* (2014)) highlighted several limitations. First, many algorithms either do not handle repeats by design or scale poorly in their presence, since the number of pairwise local alignments grows quadratically as a function of a repeat’s copy number. In addition, many algorithms use a repeat database to mask high-frequency repeats. However, these databases are usually incomplete and even a small amount of unmasked repeats may severely degrade alignment performance. Second, the number of pairwise alignments is quadratic in the number of genomes, and only a few existing approaches could handle more than ten fruit fly genomes (Earl *et al.*, 2014). Therefore, these approaches are ill-suited for large numbers of long and complex genomes, such as mammalian genomes in general and the recently assembled 16 strains of mice (Lilue *et al.*, 2018) in particular.

Alternatively, anchor based strategies can be applied to decompose genomes into locally collinear blocks (Darling *et al.*, 2004). These are blocks that are free from non-linear rearrangements, such as inversions or transpositions. Once such blocks are identified, they can independently be globally aligned (Darling *et al.*, 2004; Dewey, 2007; Paten *et al.*, 2008; Darling *et al.*, 2010; Minkin *et al.*, 2013a). The problem of constructing blocks from anchors is known as the chaining problem which had been extensively studied in the past (Myers, 1995; Abouelhoda and Ohlebusch, 2005; Ohlebusch and Abouelhoda, 2006). All of the methods applicable to datasets consisting of multiple genomes are heuristic since the exact algorithms depend exponentially on the number of genomes. Such strategies are generally better at scaling to handle repeats and multiple genomes since they do not rely on the computationally expensive pairwise alignment.

A promising strategy to find collinear blocks is based on the compacted de Bruijn graph (Raphael *et al.*, 2004; Pham and Pevzner, 2010; Minkin *et al.*, 2013b) widely used in genome assembly. Though these approaches do not work well for divergent genomes, they remain fairly accurate for closely related genomes. For example, Sibelia (Minkin *et al.*, 2013b) can handle repeats and works for many bacterial genomes; unfortunately, it does not scale to longer genomes. However, the last three years has seen a breakthrough in the efficiency of de Bruijn graph construction algorithms (Marcus *et al.*, 2014; Chikhi *et al.*, 2016; Baier *et al.*, 2016; Minkin *et al.*, 2017). The latest methods can construct the graph for tens of mammalian genomes in minutes rather than weeks. We therefore believe the de Bruijn graph approach holds the most potential for enabling scalable multiple whole-genome alignment of closely related genomes.

In this paper, we describe an algorithm SibeliaZ-LCB for identifying collinear blocks in closely related genomes. SibeliaZ-LCB is suitable for detecting homologous sequences which have evolutionary distance to the most recent common ancestor (MRCA) of at most 0.09 substitutions per site. SibeliaZ-LCB is based on the analysis of the compacted de Bruijn graph and uses a graph model of collinear blocks similar to the “most frequent paths” introduced by Cleary *et al.* (2017). This allows it to maintain a simple, static, data structure, which scales easily and allows simple parallelization. Thus, SibeliaZ-LCB overcomes a bottleneck of previous state-of-the-art de Bruijn graph-based approaches (Pham and Pevzner, 2010; Minkin *et al.*, 2013a), which relied on a dynamic data structure which was expensive to update. Further, we extend SibeliaZ-LCB into a multiple whole-genome aligner called SibeliaZ. SibeliaZ works by first constructing the compacted de Bruijn graph using our previously published TwoPaCo tool (Minkin *et al.*, 2017), then finding locally collinear blocks using SibeliaZ-LCB, and finally, running a multiple-sequence aligner (spoa, Vaser *et al.* (2017)) on each of the found blocks. To demonstrate the scalability and accuracy of our method, we compute the multiple whole-genome alignment for a collection of recently assembled strains of mice. We also test how our method works under different conditions, including various levels of divergence between genomes and different parameter settings. Our software is freely available at https://github.com/medvedevgroup/SibeliaZ/.

## 2 Results

### 2.1 Algorithm overview

As described in the introduction, the major algorithmic innovation of this paper is the SibeliaZ-LCB algorithm. SibeliaZ-LCB takes as input a de Bruijn graph built on a collection of assembled genomes. An assembled genome is itself a set of contig sequences. SibeliaZ-LCB identifies and outputs all non-overlapping blocks of homologous subsequences of the input genomes. A block can be composed of two or more sequences from one or more genomes. In this subsection, we will give a high level overview of SibeliaZ-LCB, leaving the more formal and detailed version for the Methods.

SibeliaZ-LCB relies heavily on the de Bruijn graph of the genomes. In this graph, the vertices correspond to the *k*-mers (substrings of fixed length *k*) of the input. A *k*-mer that appears multiple times in the input is represented using just one node. Then, *k*-mers that appear consecutively in some input sequence are connected by an edge from the left one to the right one (see Figure 1a for an example). This way, each genome corresponds to a path in the graph that hops from *k*-mer to *k*-mer using the edges.

In this graph, two homologous sequences form what is called a chain: an interleaving sequence of parallel edges, which correspond to identical sequences, and “bubbles”, which correspond to small variations like single nucleotide variants or indels. However, the concept of a chain is difficult to extend to more than two homologous sequences because the tangled pattern in the graph is difficult to precisely define (see Figure 1b for an example).

**Figure 1:**
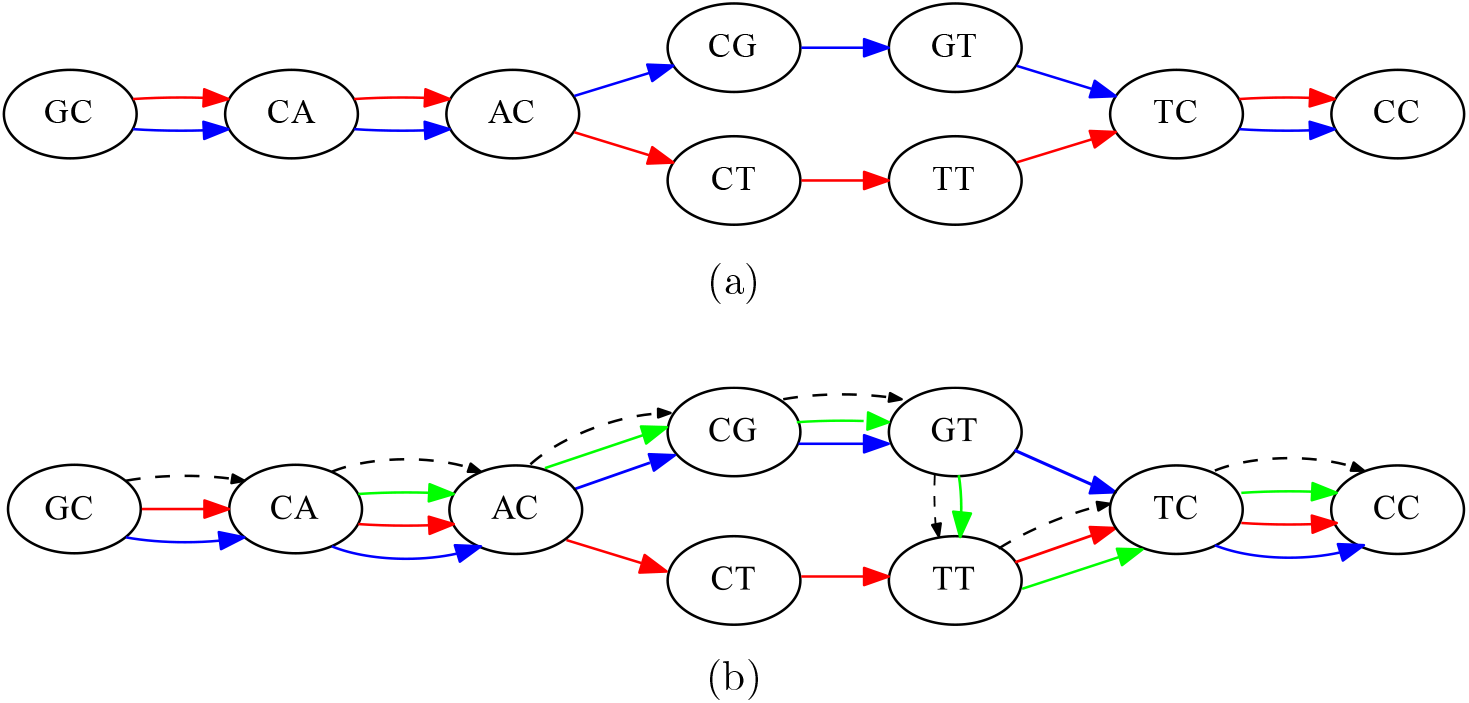
The de Bruijn graph and an example of a collinear block. (a) The graph built from strings “GCACGTCC” and “GCACTTCC”, with *k* = 2. The two strings are reflected by the blue and red walks, respectively. This is an example of a collinear block from two walks. There are four bubbles. The bubble formed by vertices “AC” and “TC” describes a substitution within the block, while three other bubbles are formed by parallel edges. The blue and red walks form a chain of four consecutive bubbles. (b) An example of more complex block, where we have added a third sequence “CACGTTCC” (green) to the input. We can no longer describe the block as a chain of bubbles, as they overlap to form tangled structures. Instead, we consider the path in the graph (dashed black) that shares many vertices with the three collinear walks. This carrying path shares many vertices with the three extant walks, and each walk forms its own chain with it. The task of finding good collinear blocks can then be framed as finding carrying paths that form good chains with the genomic walks.

To address this challenge, we introduce the idea that each block has a “carrying path” in the de Bruijn graph that holds the block together. The basic idea is that the homologous sequences forming the block have a lot of shared *k*-mers and their corresponding genomic paths go through nearly the same vertices. A carrying path is then a path that goes through the most frequently visited vertices, loosely similar to the notion of a consensus sequence from alignment. Each genomic path from the block then forms a chain with this carrying path (see Figure 1b for an example).

We do not know the carrying paths in advance but we can use them as a guiding mechanism to find blocks. We start with an arbitrary edge *e* in the graph and all other genomic paths that form bubbles with *e*. We make *e* the starting point of a carrying path and use it along with the other genomic paths to initiate the collection of sequences making up the block corresponding to this carrying path. To extend the carrying path, we look at the edges extending the genomic paths in the current block and take the most common one. The data structures maintaining the genomic paths in the block and the carrying path are then updated and the extension procedure repeats. Figure 2 shows an example of running this algorithm. We continue this process until the scoring function that describes how well a carrying path holds the block together falls below zero. At that point, we consider the possibility that we may have over-extended the block and should have instead ended it earlier. To do this, we look at all the intermediate blocks we had created during the extension process and output the one that has the highest score. Once a block is output, we output all its constituent edges as used so that they are not chosen as part of a future block.

**Figure 2:**
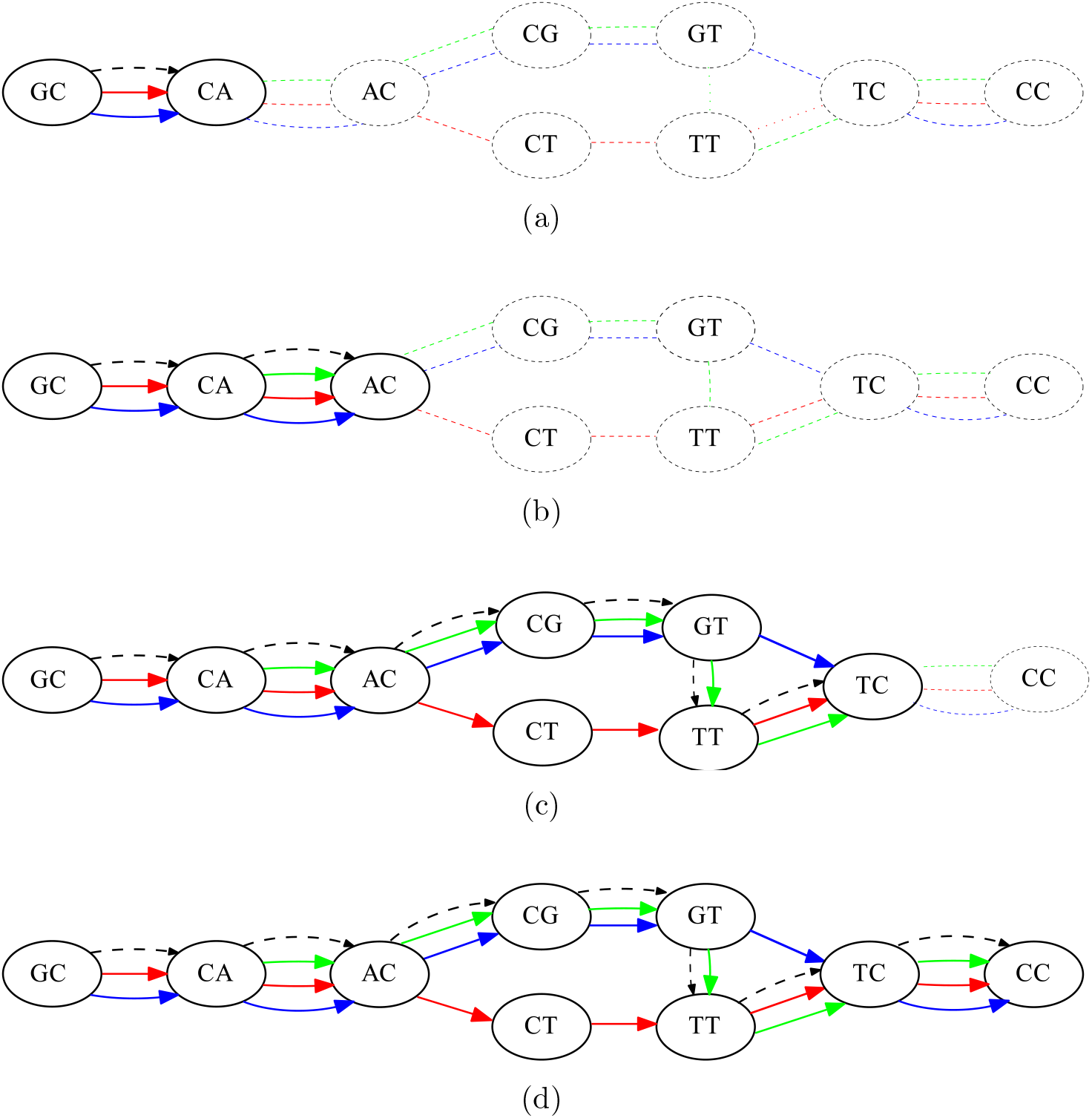
An example of running Algorithm 1 on the graph from Figure 1b, starting from edge GC → CC as the seed. Each subfigure shows the content of the collinear block *P* and the carrying path. The collinear walks are solid, the carrying path is dashed, and the rest of the graph is dotted. Subfigure (a) shows the state of these variables after the initialization; subfigures (b-d) show the state after the completion of each phase.

In this way, SibeliaZ-LCB finds a single block. Afterwards, we try to find another block by starting from another arbitrary edge. This process continues until all the edges in the graph are either used or had been tried as potential starters for a carrying path.

### 2.2 Datasets, tools, and evaluation metrics

Evaluation of multiple whole-genome aligners is a challenging problem in its own right and we therefore chose to use the practices outlined in the Alignathon (Earl *et al.*, 2014) competition as a starting point. They present several approaches to assess the quality of a multiple whole-genome alignment. Ideally it is best to compare an alignment against a manually curated gold standard; unfortunately, such a gold standard does not exist. We therefore chose to focus our evaluation on real data.

We evaluated the ability of SibeliaZ to align real genomes by running it on several datasets consisting of varying number of mice genomes. We retrieved 16 mice genomes available at GenBank (Benson *et al.*, 2017) and labeled as having a “chromosome” level of assembly. They consist of the mouse reference genome and 15 different strains assembled as part of a recent study (Lilue *et al.*, 2018) (Table S1). The genomes vary in size from 2.6 to 2.8 Gbp and the number of scaffolds (between 2,977 and 7,154, except for the reference, which has 377). We constructed 4 datasets of increasing size to test the scalability of the pipelines with respect to the number of input genomes. The datasets contain genomes 1-2, 1-4, 1-8 and 1-16 from Table S1, with the genome 1 being the reference genome.

To measure accuracy, we used several ground truth alignments (to be described) and employed the metrics of precision and recall used in the Alignathon and implemented by the mafTools package (Earl *et al.*, 2014). For these metrics, alignment is viewed as an equivalence relation. We say that two positions in the input genomes are equivalent if they originate from the same position in the genome of their recent common ancestor. We denote by *H* the set of all equivalent position pairs, participating in the “true” alignment. Let *A* denote the relation produced by an alignment algorithm. The accuracy of the alignment is then given by recall(*A*) = 1 − |*H* \ *A*|*/*|*H*| and precision(*A*) = 1 − |*A* \ *H*|*/*|*A*|, where \ denotes set difference.

To evaluate recall, we compared our results against annotations of protein-coding genes. We retrieved all pairs of homologous protein-coding gene sequences from Ensembl and then computed pairwise global alignments between them using LAGAN (Brudno *et al.*, 2003). The alignment contains both orthologous and paralogous genes, though most of the paralogous pairs come from the well-annotated mouse reference genome. We removed any pairs of paralogous genes with overlapping coordinates, as these were likely mis-annotations, as confirmed by Ensembl helpdesk (Perry, 2018). We made these filtered alignments as well as the alignments produced by SibeliaZ available for public download from our GitHub repository (see Section 4.4 for the links).

We define the nucleotide identity of an alignment as the number of matched nucleotides divided by the length of an alignment, including gaps. The distribution of nucleotide identities as well as the coverage of the annotation is shown in Figure S1. In our analysis, we binned pairs of genes according to their nucleotide identity.

Since protein-coding genes only compromise a small portion of the genome, we also computed all-against-all pairwise local alignments between chromosomes 1 of genomes 1-2 and 1-4 using LASTZ (Harris, 2007), a reliable local aligner known for its accuracy. We only computed alignments between chromosomes of different genomes, i.e. did not include self-alignments, which excludes duplications such as paralogous genes from the alignment. We used default settings of LASTZ except that we made it output alignments of nucleotide identity at least 90%. We then evaluated the recall and precision of our alignments but restricted our alignments to the sequences of chromosome 1. We then treated the LASTZ alignments as the ground truth. The LASTZ alignments are available for download from our repository’s supplemental data section. Note that because the alignment is represented as a set of positions pairs, it is possible to evaluate a multiple whole-genome alignment using pairwise local alignments.

To measure precision, we use the LASTZ alignments on chromosome 1. However, it is computationally prohibitive to compute such alignments with LASTZ for the whole genome. We therefore also use an indirect way to assess precision for the whole genome. For each column in the alignment we calculate the average number of nucleotide differences (Tajima, 1989). In an alignment of highly similar genomes that has high precision, we expect these numbers to be low (close to 0) for most of the columns in the alignment. Otherwise, it would suggest presence of unreliable poorly aligned blocks in the alignment. Formally, given a column *c* of a multiple whole-genome alignment with *c_i_* being its *i*-th element, average number of nucleotide differences is given by 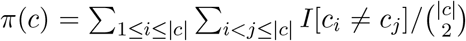. The variable *I*[*c*_*i*_ ≠ *c*_*j*_] is equal to 1 if both *c_i_* and *c_j_* are different valid DNA characters and 0 otherwise; |*c*| is the number of rows in the column *c*.

We benchmarked the performance of SibeliaZ against Progressive Cactus (Armstrong *et al.*, 2019), an aligner based on analysis of the Cactus graphs Paten *et al.* (2011b) built from pairwise alignments. We also attempted to run Sibelia (Minkin *et al.*, 2013b) (a predecessor of SibeliaZ) and MultiZ+TBA (Blanchette *et al.*, 2004), but these could run to completion within a week on even a single mouse genome. Other multiple aligners (Dubchak *et al.*, 2009; Darling *et al.*, 2010; Angiuoli and Salzberg, 2011) benchmarked in the Alignathon could not handle a dataset of 20 flies and hence are unlikely to scale to a mammalian dataset. We also chose to not run Mercator (Dewey, 2007) since it requires a set of gene exons as input and hence solves a different problem: in this paper we focus on computing the whole-genome alignment directly from the nucleotide sequences without using external information. Further details about parameters, versions, and hardware are in Supplementary Note 2.

### 2.3 Running time and memory

The running times of SibeliaZ and Cactus are shown on Figure 3 (Table S2 contains the raw values). On the dataset consisting of 2 mice, SibeliaZ is more than 10 times faster than Cactus, while on 4 mice SibeliaZ is more than 20 times faster. On the datasets with 8 and 16 mice, SibeliaZ completed in under 7 and 16 hours, respectively, while Cactus did not finish (we terminated it after a week). For SibeliaZ, we note that the global alignment with spoa takes 44-73% of the running time, and, for some applications (e.g. rearrangement analysis), this step may be further omitted to save time. Memory is shown in Table S2. When it is able to complete, Cactus has better memory performance than SibeliaZ; however, both tools require memory that is well within the range of most modern servers but outside the range of personal computers.

**Figure 3:**
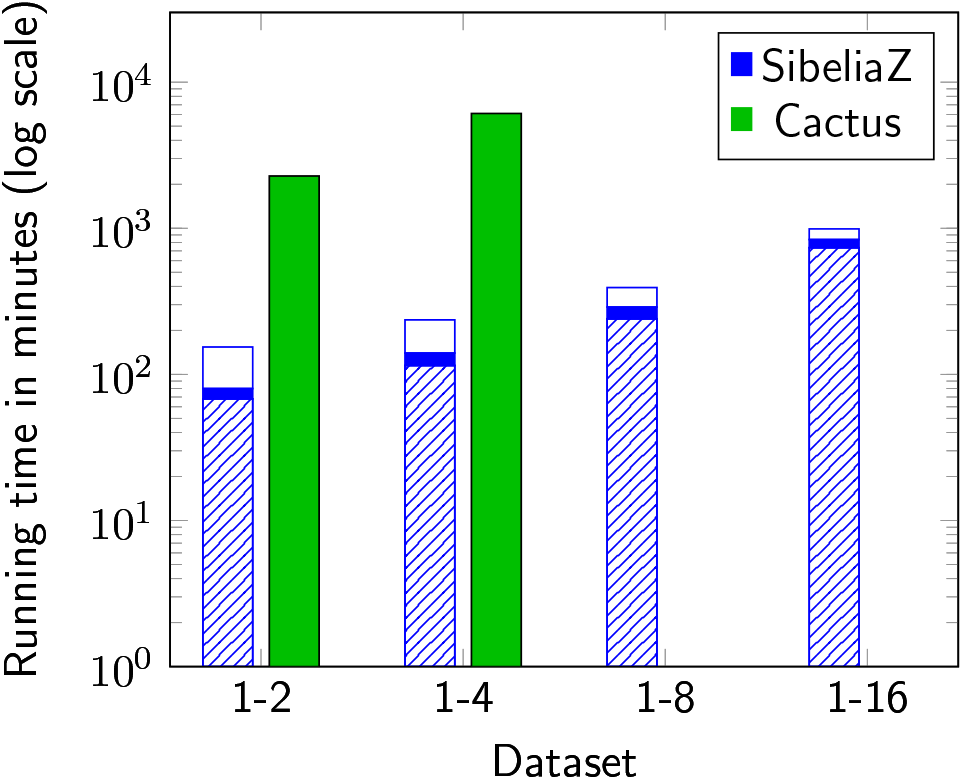
Running times of the different pipelines on the mice datasets (on a log scale). Each bar corresponds to a pipeline. The bar of SibeliaZ is split according to its components: spoa (hatch fill), TwoPaCo (solid fill), and SibeliaZ-LCB (empty fill). Cactus is not shown on datasets 1-8 and 1-16 because it did not complete. We used 32 threads for each experiment.

### 2.4 Accuracy

In Tables 1 and 2, we show the properties of the alignments found by SibeliaZ and Cactus. To compute recall, we only used nucleotides from gene pairs having at least 90% identity in the annotation. For the datasets where Cactus was able to complete, SibeliaZ had and similar recall on orthologous pairs. We did not evaluate the results on paralogs by Cactus since it heuristically filters out paralogous alignments (Armstrong *et al.*, 2019) as a part of its pipeline. SibeliaZ’s recall decreases only slightly going up to the whole 16 mice dataset, indicating that the recall scales with the number of genomes.

**Table 1:** Number of blocks and coverage by the multiple whole-genome alignments computed by SibeliaZ and Cactus from the mice datasets.

| Dataset | N. of blocks, SibeliaZ | N. of blocks, Cactus | Coverage, SibeliaZ | Coverage, Cactus |
| --- | --- | --- | --- | --- |
| 1-2 | 2,083,258 | 4,228,063 | 0.88 | 0.85 |
| 1-4 | 2,739,821 | 6,133,662 | 0.86 | 0.84 |
| 1-8 | 3,179,619 | - | 0.89 | - |
| 1-16 | 4,507,109 | - | 0.88 | - |

**Table 2:** Recall of the orthologous and paralogous basepairs by the multiple whole-genome alignments computed by SibeliaZ and Cactus from the mice datasets, using Ensembl gene annotation as the ground truth. Recall of paralogs by Cactus is not included (see text).

| Dataset | Ort. nt. pairs, SibeliaZ | Ort. nt. pairs, Cactus | Par. nt. pairs, SibeliaZ |
| --- | --- | --- | --- |
| 1-2 | 0.99 | 0.99 | 0.89 |
| 1-4 | 0.98 | 0.98 | 0.89 |
| 1-8 | 0.98 | - | 0.84 |
| 1-16 | 0.98 | - | 0.83 |

We also measured coverage, which is the percent of the genome sequence that is included in the alignment. The coverage of both tools is roughly the same, but SibeliaZ has only about half the blocks. The different amounts of blocks produced by the tools are likely to be a result of the different approaches to the formatting of the output. Representation of multiple whole-genome alignment is ambiguous and the same alignment can be formatted in different but mathematically equivalent forms varying by the number blocks.

We further investigate how the recall behaved as a function of nucleotide identity, for the two- and four-mice dataset (Figure 4). As expected, recall decreases with nucleotide identity, though SibeliaZ’s recall remains above 90% for nucleotides from similar (80-100% identity) orthologous genes. Cactus has slightly better recall in orthologous genes of lower identity on the two-mice dataset. We note that the gene annotation was constructed (Lilue *et al.*, 2018) using an alignment produced by Cactus which was further processed by annotation software CAT (Fiddes *et al.*, 2018). This fact might give Cactus a slight advantage in this comparison and explain why Cactus has slightly better recall. Recall on orthologous gene pairs remains consistent in both two- and for-mice datasets for both datasets.

**Figure 4:**
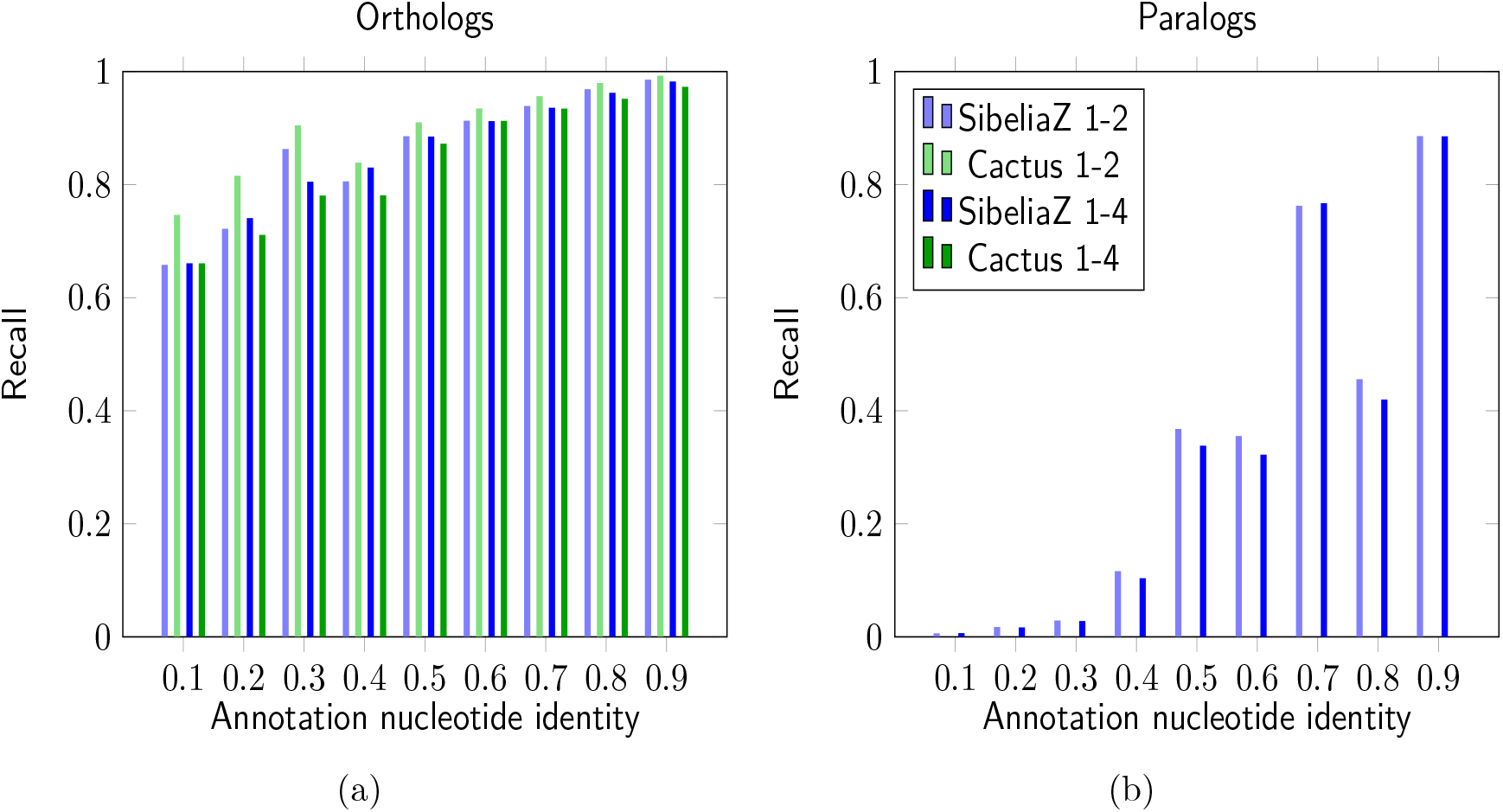
Recall of orthologous (a) and paralogous (b) nucleotide pairs, on the 1-2 and 1-4 mouse datasets. Nucleotide pairs are binned according the nucleotide identity of their respespective genes in the annotation. Recall of paralogs by Cactus is not shown (see text).

At the same time, we observed a much less consistent picture for paralogous pairs of genes. For example, SibeliaZ was able to recover nearly 90% of the paralogous basepairs belonging to gene pairs of nucleotide identity of 90%, but found less than 45% of the basepairs of gene pairs of 80% identity.

The results of the precision and recall measured with respect to LASTZ alignments are shown in Table S3. On the dataset consisting of two genomes, Cactus had slightly higher recall of 0.97 versus 0.95 of SibeliaZ. On the other hand, SibeliaZ had better precision: 0.93 against 0.89 of Cactus. With the four genomes, SibeliaZ maintained its recall of 0.95 while recall of Cactus dropped to 0.92. On this dataset SibeliaZ also had higher precision: 0.96 and 0.90, respectively. Overall, these numbers show that the alignment accuracy of SibeliaZ and Cactus is similar.

Finally, since we could not evaluate genome-wide precision, we use the proxy of the average number of nucleotide differences for the alignment columns (Figure S2). SibeliaZ’s alignment has a high degree of similarity: more than 95% of the alignment columns have *π*(*c*) ≤ 0.1, which is what we would expect from aligning closely related genomes. Cactus has slightly lower percentage of highly similar columns, which may simply indicate that it finds more blocks with higher divergence.

We note that the results in this Section evaluate the accuracy of SibeliaZ-LCB and spoa simultaneously; however, since SibeliaZ is targeted at closely related genomes, we observed that the global alignment procedure has a negligible effect on accuracy (data not shown). This is due to the fact that the global alignment of similar sequences is likely to be unambiguous at homologous nucleotides and robust with respect to different algorithms and their parameters.

### 2.5 Results on simulated data

In addition to the real data, we measured performance of different whole-genome aligners on a larger simulated dataset with small genomic divergence, called “primates” in (Earl *et al.*, 2014). In this dataset the distance from the root to the leaves in the phylogenetic tree is equal to 0.02 substitutions per site. The dataset has four genomes, with four chromosomes each, and each genome is approximately 185 Mbp in size. We did not use the other simulated dataset in (Earl *et al.*, 2014) since its divergence of around 0.4 substitutions per site is outside of the target range of SibeliaZ.

On this dataset, SibeliaZ pipeline was 20 times faster than Cactus and consumed 2.5 times less memory: SibeliaZ finished in 18 minutes using 7 GBs of memory, while Cactus took 363 minutes to finish and used 18 GBs of memory. Sibelia and MultiZ could not finish on the “primates” dataset in a week. Table 3 demonstrate the recall and precision values for the alignments produced by SibeliaZ and Cactus on this dataset. SibeliaZ showed recall of 95% and precision of 92%, while Cactus had 98% recall and 95% precision. We note that according to Earl *et al.* (2014) the precision values calculated using this dataset can be considered lower bounds due to the features of the simulation process. Particularly, the ground truth for this dataset is likely to miss some repetitive alignments, hence we believe that the lower precision values of SibeliaZ may be due to this reason.

**Table 3:** Recall and precision of the alignments computed by Cactus and SibeliaZ on the “primates” dataset from the Alignathon.

| Program | Recall | Precision |
| --- | --- | --- |
| SibeliaZ | 0.95 | 0.92 |
| Cactus | 0.98 | 0.95 |

### 2.6 Gene families

We wanted to further understand SibeliaZ’s ability to recall homologous nucleotides from large gene families. Aligning genes having many copies is a challenging task since they generate a tangled de Bruijn graph. To investigate, we took each pair of genes in the two-mice dataset that have greater than 90% nucleotide identity. We then identify any other homologous genes that had a nucleotide identity of at least 90% to one of the genes in the pair. We refer to the number of such genes as the inferred family size of the gene pair, which roughly corresponds to the gene family size in the biological sense. Figure S3 then shows the recall of nucleotide pairs with respect to the inferred family size of their respective genes. The recall shows a lot of variance with respect to the inferred family size but does exhibit a general trend of decreasing with increasing family size. The largest bin (with inferred family size of 58) corresponds to a single large gene family on the Y chromosome (*PTHR19368*) and actually has relatively high recall.

This experiment shows that finding all copies of even very similar homologous sequences within long genomes can be a challenging task. Moreover, the high variance we observe indicates that this challenge cannot be reduced to a single factor like family size. A manual inspection of false negatives suggests that the drop in recall may be due to complex substructures of unannotated repeats forming tangled graph structures.

### 2.7 Effect of parameters and sequence divergence

SibeliaZ-LCB has four primary parameters that affect its performance. The most critical dependence is on the size of a *k*-mer (i.e. *k*) and the maximum allowed length of a bubble *b*. For a given sequence divergence, the distance between shared *k*-mers forming bubbles in homologous regions increases with *k*. At the same time, the maximum allowed length of a bubble is *b*. If the distance exceeds *b*, then SibeliaZ may fail to uncover such regions and result in lower recall. To avoid this situation, we can either decrease *k* or increase *b*. Decreasing *k* is desirable up to a point, but when *k* becomes too low, the de Bruijn graph becomes convoluted and our algorithm becomes more time and memory consuming. Increasing *b* can also be done but simultaneously increases the allowable gap length, leading to decreased precision.

Over-alignment is the problem of combining non-homologous sequences in a single block, which is closely related to low precision (Schwartz and Pachter, 2007). In our case, one can control over-alignment by looking at the *π*(*c*) scores, as we have done in our analysis (Figure S2). A higher score indicates that more divergent sequences are included in a block. If the divergence is deemed too high by the user, it is recommended to reduce *b*.

To investigate this complex interplay between *k* and *b* and its relationship to sequence divergence, we used simulations (Supplementary Note 3) to measure recall (Figure S4) and precision (Figure S5) under various combinations. As predicted, recall increases with decreasing *k* and with increasing *b*, and precision decreases with increasing *b*. We note though that the precision varies only a little and remains high. Based on these analyses, we recommend two values of *k* for practical usage. For less complex organisms (e.g. bacteria), we recommend *k* = 15, since it yields the highest recall. This value is impractical for complex organisms (e.g. mammals) due to runtime, so we recommend setting *k* = 25 in those cases as it provides a reasonable trade-off between accuracy and required computational resources (we used this for our mice datasets). For the value of *b*, we observed that increasing *b* lowers the precision at only higher values. Hence, we recommend *b* = 200 as the default in all cases, as it led to high recall across all tested ranges of *k* on our simulated data without lowering precision.

To test the level of divergence which SibeliaZ-LCB can tolerate, we took the default values of *k* = 15 or 25 and *b* = 200 and plotted the precision vs. recall curve as a function of the root-to-leaf divergence of the dataset (Figure S6). We see that for *k* = 25 the recall deteriorates significantly for datasets having a root-to-leaf evolutionary distance of more than 0.09 substitutions per site. Based on this, we recommend that for large datasets SibeliaZ-LCB be only used for detecting homologs with an evolutionary distance to the MRCA of at most 0.09 substitutions per site.

The other two parameters that can effect SibeliaZ-LCB’s performance are the minimum size of a locally collinear block *m* and the abundance pruning parameter *a*. These parameters should be set according to the type of data and its intended use. The parameter *m* controls the fragmentation of the alignment and the coverage — higher *m* results in longer blocks spanning less of the genomes, since short blocks are not reported. We recommend the parameter *m* to be set to the length of the shortest homologous sequence of interest to the downstream analysis. We set *m* = 50 as a default, since this is smaller than 93.1% of the known mice exons (Sakharkar *et al.*, 2005) and, more generally, we do not expect most applications to be interested in much blocks shorter than 50nt. In the case that a user is interested in larger homologous units, they can increase *m* together with *b*. Alternatively, they can use either synteny block generation or alignment chaining algorithms for post-processing the alignments produced by SibeliaZ (see Supplementary Note 1 for relevant references).

The abundance pruning parameter *a* is a filtering parameter for *k*-mers whose abundance is above *a*. Such *k*-mers are still considered by SibeliaZ-LCB, but to a smaller extent, resulting in reduced recall in regions with such *k*-mers. We recommend setting *a* as high as the compute resources allow, keeping in mind that homologous blocks with multiplicity higher than *a* are possibly not going to be captured. For the mice dataset, we used *a* = 150.

## 3 Discussion

In this paper, we presented a whole-genome alignment pipeline SibeliaZ based on an algorithm for identifying locally collinear blocks. The algorithm analyses the compacted de Bruijn graph and jointly reconstructs the path corresponding to a collinear block and identifies the induced collinear walks. We assume that the collinear walks share many vertices with this carrying path and form chains of bubbles. Each carrying path and the induced block is found greedily, using a scoring function that measures how close it is to all the sequences in the block. We then globally align the collinear blocks to generate the whole-genome alignment.

SibeliaZ builds on the ideas laid down in DRIMM-Synteny (Pham and Pevzner, 2010) and Sibelia (Pham and Pevzner, 2010) that used variants of the de Bruijn graphs for finding synteny blocks (we elaborate on the connection between the whole genome alignment and synteny reconstruction in Supplementary Note 1). Sibelia did not scale beyond bacterial genomes due to its slow graph construction algorithm and the fact that it continuously had to modify the graph. TwoPaCo (Minkin *et al.*, 2017) addressed the former issue and we use it as a standalone module in SibeliaZ. The latter issue was addressed in this paper by SibeliaZ-LCB, which achieves its speed in part because it keeps the underlying graph static.

The main strength of our approach is speed — we achieve drastic speedups compared to the state-of-the-art Progressive Cactus aligner (Armstrong *et al.*, 2019) while retaining comparable accuracy. Using a single machine, on 16 mice genomes, SibeliaZ runs in under 16 hours, while Progressive Cactus is not able to complete for even 8 mice genomes, within seven days. We note that it is possible for Cactus to construct larger alignments by utilizing a distributed computer cluster (Armstrong *et al.*, 2019). In our study, we concentrated on improving scalability of the whole-genome alignment when only a single machine is available. In the future we hope to develop a version of SibeliaZ that will work in the distributed setting as well. Overall, SibeliaZ is the only tool available that can scale to many long, closely related genomes on a single machine.

The biggest limitation of our approach is the limited tolerance to the divergence of input sequences. As suggested by the results on simulated bacterial data, SibeliaZ works best for aligning genomes having an evolutionary distance to the most recent common ancestor of at most 0.09 substitutions per site. Aligning more divergent genomes with SibeliaZ is still possible but it will result in smaller recall; for such datasets, Cactus remains a better option. In the future, we hope to address this limitation by employing techniques like postprocessing of the output with more sensitive homology finders.

If the alignments themselves are not needed, SibeliaZ-LCB can be run alone (without spoa) to construct the collinear blocks. This is most useful in applications stemming from studies of genome rearrangements, which can be applied to study breakpoint reuse (Pevzner and Tesler, 2003b), ancestral genome reconstruction (Kim *et al.*, 2017) and phylogenies (Luo *et al.*, 2012). Locally collinear blocks are also a required input for scaffolding tools using multiple reference genomes (Kim *et al.*, 2013; Kolmogorov *et al.*, 2014; Chen *et al.*, 2016; Aganezov and Alekseyev, 2016). For such applications, the output of SibeliaZ-LCB can be used either directly or after postprocessing by a synteny block generator (Pham and Pevzner, 2010; Proost *et al.*, 2011).

There are several remaining open questions of interest. A formal analysis of SibeliaZ-LCB’s runtime is relevant, but doing it in a useful way is a challenge. The worst-case running time does not reflect the actual one; moreover, we observed that the actual one depends on multi-thread synchronization, which is challenging to model. However, it would be interesting if such a time analysis can be performed parametrized by the crucial properties of the structure of the input. We also did not investigate how close to an optimal solution our greedy heuristic gets. One way to do this would be to find an optimal carrying path using exhaustive enumeration, but the search space even for a small realistic example is too big. We suspect that a polynomial time optimal solution is not possible, but the computational complexity of our problem is open.

SibeliaZ is the first multiple whole-genome aligner that can run on a single machine in reasonable time on a dataset such as the 16 mouse genomes analyzed in this paper. With ongoing initiatives like the Vertebrate Genomes Project and the insect5k, thousands of species will soon have a reference genome available, and the sequencing and assembly of various sub-species and strains will be the next logical step for many comparative genomics studies. For example, Portwood *et al.* (2018) currently holds 18 assembled maize genomes, with more to come in the recent future. Similarly, the Solanaceae Genomics Network has recently released the genomes of 13 diverse tomato accessions (https://solgenomics.net/projects/tomato13/). Analysis of such datasets is likely to be carried out in single-lab settings with limited compute resources, rather than at large computing centers like EMBL or NCBI. SibeliaZ makes a significant leap towards enabling such studies.

## 4 Methods

### 4.1 Preliminaries

First, we will define the de Bruijn graph and related objects. Given a positive integer *k* and a string *s*, we define a multigraph *G*(*s, k*) as the *de Bruijn graph* of *s*. The vertex set consists of all substrings of *s* of length *k*, called *k*-mers. For each substring *x* of length *k* + 1 in *s*, we add a directed edge from *u* to *v*, where *u* is the prefix of *x* of length *k* and *v* the suffix of *x* of length *k*.

Each occurrence of a (*k* + 1)-mer yields a unique edge, and every edge corresponds to a unique location in the input. Two edges are *parallel* if they are oriented in the same direction and have the same endpoints. Note that two edges are parallel if and only if they were generated by the same (*k* + 1)-mer. This way, we use the notion of parallel edges to refer to a set of identical (*k* + 1)-mers in the input strings. Parallel edges are not considered identical. The de Bruijn graph can also be constructed from a set of sequences *S* = {*s*_1_*, …, s_n_*}. This graph is the union of the graphs constructed from the individual strings: *G*(*S, k*) = ∪_1≤*i*≤*n*_ *G*(*s_i_, k*). See Figure 1 for an example. The set of all edges in a graph *G* is denoted by *E*(*G*). We write (*u, v*) to denote a edge from vertex *u* to *v*. A *walk p* is a sequence of edges ((*v*_1_*, v*_2_), (*v*_2_*, v*_3_)*, …,* (*v*_|*p*|−1_*, v*_|*p*|_)) where each edge (*v_i_, v_i_*_+1_) belongs to *E*(*G*). The length of the walk *p*, denoted by |*p*|, is the number of edges it contains. The last edge of a walk *p* is denoted by end(*p*). A path is a walk that visits each vertex at most once.

In a de Bruijn graph, a given edge *x* was generated by a (*k* + 1)-mer starting at some position *j* of some string *s_i_*. To retrieve the position *j* of the (*k* + 1)-mer that generated edge *x*, we define function pos(*x*) = *j*. We use the function *pos* to map edges of the graph back to positions of the *k*-mers that generated them. For an edge *x*, its successor, denoted by next(*x*), is an edge *y* such that both *x* and *y* are generated by the same string and pos(*y*) = pos(*x*) + 1. Note that a successor does not always exist.

A walk *p* = (*x*_1_*, …, x*_|*p*|_) is *genomic* if nex*t*(*x_i_*) = *x_i_*_+1_ for 1 ≤ *i* ≤ |*p*| − 1. In other words, a walk is genomic if it was generated by a substring in the input. The *b-extension* of a genomic walk *p* is the longest genomic walk *q* = (*y*_1_*, …, y*|*q*|) such that *y*_1_ = next(end(*p*)) and |*q*| ≤ *b*. The *b*-extension of a walk *p* is uniquely defined and usually has length *b*, unless *p* was generated by a substring close to an end of an input string. Intuitively, *b*-extension defines edges lying ahead of a walk. As our algorithm works in the seed-and-extend manner, we later use *b*-extensions to find the most appropriate extension of the current block. The concatenation of two walks *x* and *y* is a walk (if it exists) *xy* consisting of edges of *x* followed by edges of *y*. We use the concatenation operation to extend the genomic walks constituting locally-collinear blocks using appropriate *b*-extensions.

### 4.2 Problem formulation

In this section, we will define the collinear block reconstruction problem in de Bruijn graphs. A *collinear block* is a set of edge-disjoint genomic walks with length at least *m*, where *m* is a parameter. We call walks constituting a collinear block *collinear walks*. In order to quantify how well collinear walks correspond to homologous sequences, we will define a *collinearity score* of a collinear block. Our problem will then be to find a set of collinear blocks that are pairwise edge-disjoint and have the largest score.

We capture the pattern of two homologous collinear walks through the concept of chains and bubbles. A *bubble* is a subgraph corresponding to a possible mutation flanked by equal sequences. Formally, a pair of walks *x* and *y* form a bubble (*x, y*) if all of the following holds: (1) *x* and *y* have common starting and ending vertices; (2) *x* and *y* have no common vertices except the starting and ending ones; and (3) |*x*| ≤ *b* and |*y*| ≤ *b*, where *b* is a parameter. A *chain c* = ((*x*_1_*, y*_1_), (*x*_2_*, y*_2_)*, …,* (*x_n_, y_n_*)) is a sequence of bubbles such that *x* = *x*_1_*x*_2_ *… x_n_* and *y* = *y*_1_*y*_2_ … *y*_*n*_ are walks in a de Bruijn graph. In other words, a chain is a series of bubbles where each bubble is a proper continuation of the previous one. Note that two parallel edges form a bubble and a chain arising from equal sequences corresponds to a series of such bubbles. This way, a chain models a pair of sequences that potentially have point mutations and indels. For an example of a bubble and a chain, see Figure 1.

The subgraph resulting from more than two collinear walks can be complex, and there are several ways of capturing it. We note that there are previous studies generalizing the idea of bubbles, see (Onodera *et al.*, 2013; Sung *et al.*, 2015; Iliopoulos *et al.*, 2016; Brankovic *et al.*, 2016; Paten *et al.*, 2017), mostly in the context of analyzing assembly graphs. We decided to follow a different approach designed specifically for capturing locally-collinear blocks.

Our approach is to give a definition that naturally leads itself to an algorithm. As homologous sequences all originate from some common ancestral sequence *s*_a_, they should have many common *k*-mers and there should be a path *p*_a_ = *G*(*s*_a_*, k*) through the graph forming a long chain with each walk *p* in the collinear block. We call such path a *carrying* one. We require the chains to be longer than *m* to avoid confusing spuriously similar sequences with true homologs. At the same time, a collinear walk may only partially form a chain with the carrying path, leaving hanging ends at the ends of the carrying path, which is undesirable since it implies that these graphs originate from dissimilar sequences. We note that compared to the previous work on chaining anchors (Myers, 1995; Abouelhoda and Ohlebusch, 2005; Ohlebusch and Abouelhoda, 2006) our definition of the block will be more relaxed. Namely, we will not require for common *k*-mers to be present in all copies of a block. In addition, many alignment methods use phylogenetic information for scoring purposes. We decided to not use it since our method targets closely-related sequences, such as strains of the same species, where the phylogeny is often unknown.

We formalize this intuition by introducing a scoring function quantifying how well a carrying path describes a collection of the collinear walks. The function rewards long chains formed by the carrying path and a collinear walk and penalizes the hanging ends. Given a carrying path *p*_a_ and a genomic walk *p*, let *q*_2_ be the longest subpath of *p*_a_ that forms a chain with *p*. Then, we can write *p*_a_ = *q*_1_*q*_2_*q*_3_. Recall that *m* is the parameter denoting the minimum length of a collinear block, and *b* is the maximum bubble size We define the score *f* (*p*_a_*, p*) as:

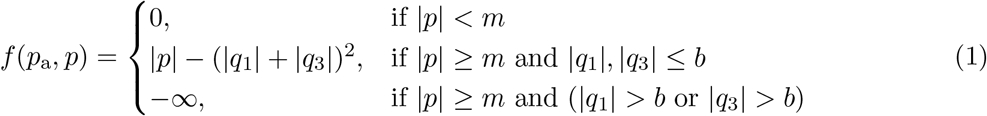

The third case forbids walks (i.e. gives them a score of −∞) where the hanging ends are too long, and the first case ignores walks (i.e gives them a score of 0) that weave through *p*_a_ but are too short. The second case gives a score that is proportional to the length of the part of *p*_a_ that forms a chain with *p*. At the same time, it reduces the score if the collinear walks leave hanging ends *q*_1_ and *q*_3_ — the parts of *p*_a_ not participating in the chain. The penalty induced by these ends is squared to remove spuriously similar sequences from from the collinear block. This form of scoring function showed better performance compared to other alternatives (data not shown). We do not penalize for discrepancy between *p* and *q*_2_ for the sake of simplicity of the scoring function and avoiding extra computation needed to calculate it. Figure 5 shows an example of computing the score.

**Figure 5:**
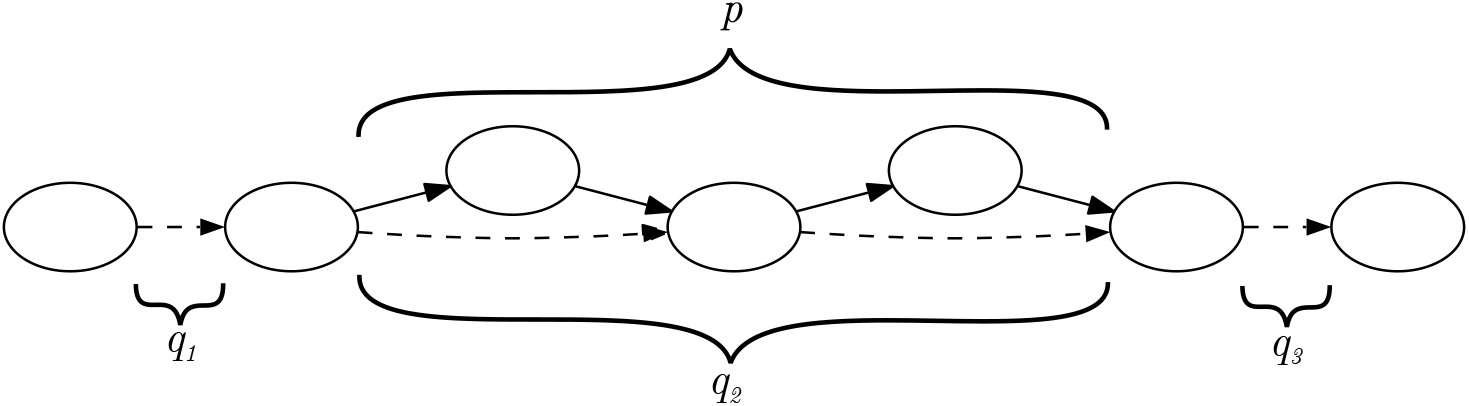
An example of computing the score of a walk *p* (solid) relative to a carrying path *p*_a_ = *q*_1_*q*_2_*q*_3_ (dashed). The path *p* forms a chain with the subpath *q*_2_ of *p*, while subpaths *q*_1_ and *q*_3_ form hanging ends. We count the length of *p* and subtract lengths of the hanging ends. Thus, the score *f* (*p*_a_*, p*) = 4 − (1 + 1)^2^ = 0.

The *collinearity score* of a collinear block is given by

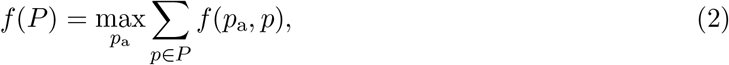

where *p*_a_ can be any path (not necessarily genomic). In other words, we are looking for a path forming longest chains with the collinear walks and thus maximizes the score. The collinear blocks reconstruction problem is to find a set of collinear blocks 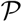 such that 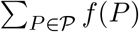 is maximum and no two walks in 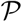 share an edge. Note that the number of collinear blocks is not known in advance. For an example of a complex collinear block in the de Bruijn graph and a carrying path capturing it, refer to Fig 1b.

### 4.3 The collinear blocks reconstruction algorithm

**Algorithm 1.**
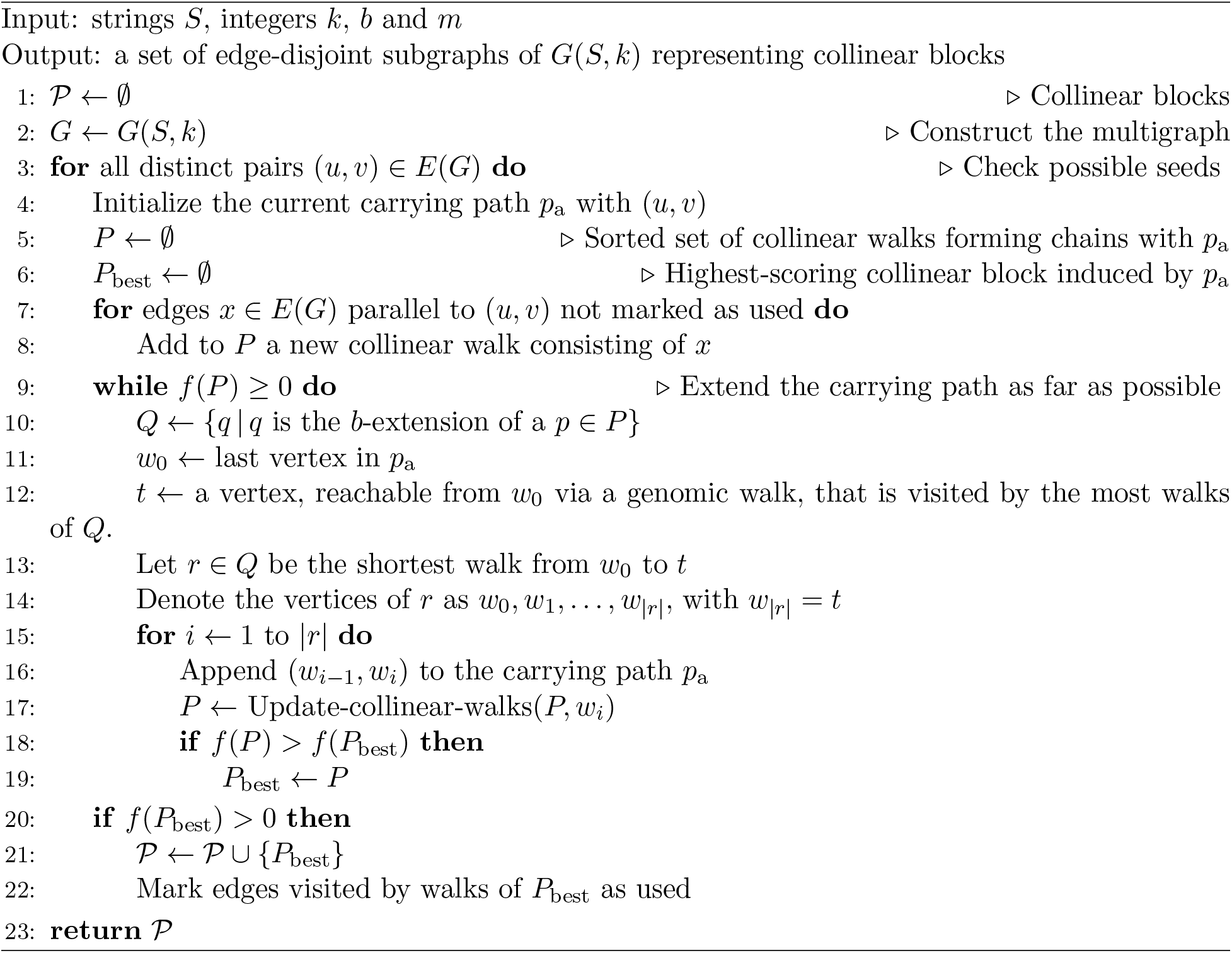
Find-collinear-blocks

**Algorithm 2.**
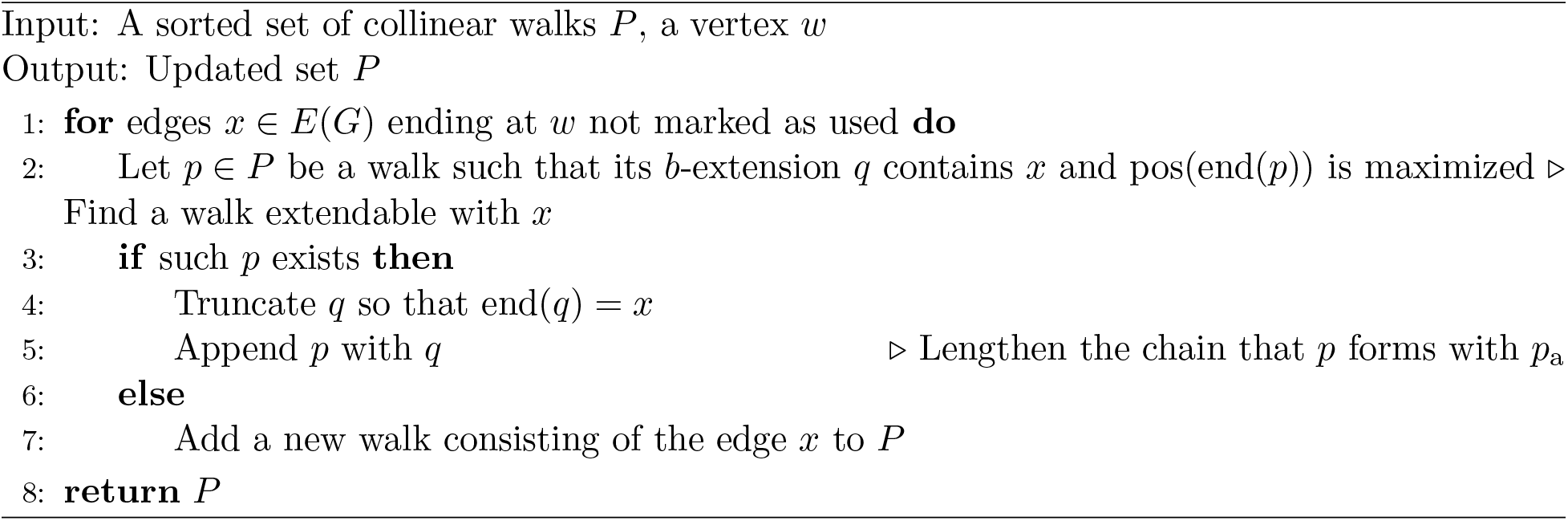
Update-collinear-walks

Our algorithm’s main pseudocode is shown in Algorithm 1 and its helper function in Algorithm 2. First, we describe the high-level strategy. The main algorithm is greedy and works in the seed-and-extend fashion. It starts with an arbitrary edge in the graph, and tries to extend it into a carrying path that induces a collinear block with the highest possible collinearity score *P*_best_. If the block has a positive score, then *P*_best_ is added to our collection of collinear blocks 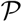. The algorithm then repeats, attempting to build a collinear block from a different edge seed. New collinear blocks cannot use edges belonging to previously discovered collinear blocks. This process continues until all possible edges are considered as seeds. The algorithm is greedy in a sense that once a block is found and added to 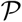, it cannot be later changed to form a more optimal global solution.

To extend a seed into a collinear block *P*, we first initialize the collinear block with a walk for each unused edge parallel to the seed (including the seed) (lines 7 to 8). These parallel edges represent the different occurences of the seed string in the input and, hence, form the initial collinear block. We then proceed in phases, where each phase is an iteration of the while loop (lines 9 to 19). During each phase, the carrying path *p*_a_ is extended using a walk *r* of length at most *b* (lines 10 to 14). Next, we try to extend each of the collinear walks in a way that forms chains with the extended *p*_a_ (lines 15 to 19). The extension of a seed into a collinear block is also a greedy process, since we only change *p*_a_ and the walks in *P* by extending them and never by changing any edges. Finally, we check that the collinearity score for our extended block is still positive — if it is, we iterate to extend it further, otherwise, we abandon our attempts at further extending the block. We then recall the highest scoring block that was achieves for this seed and save it into our final result 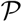 (lines 20 to 22).

To pick the walk *r* by which to extend *p*_a_, we use a greedy heuristic (lines 10 to 14). First, we pick the vertex *t* which we want to extension to reach (lines 10 to 12). We limit our search to those vertices that can be reached by a genomic walk from the end of *p*_a_ and greedily chose the one that is most often visited by the *b*-extensions of the collinear walks in *P*. Intuitively, we hope to maximize the number of collinear walks that will form longer chains with *p*_a_ after its extension and thereby boost the collinearity score. We then extend *p*_a_ using the shortest *b*-extension of the walks in *P* to reach *t*. We chose this particular heuristic because it showed superior performance comparing to other possible strategies.

Once we have selected the genomic walk *r* by which to extend *p*_a_, we must select the extensions to our collinear walks *P* that will form chains with *p*_a_*r*. This is done by the function Update-collinear-blocks (Algorithm 2). We extend the walks to match *r* by considering the vertices of *r* consecutively, one at a time. To extend to a vertex *w*, we consider all the different locations of *w* in the input (each such location is represented by an edge *x* ending at *w*). For each location, we check if it can be reached by a *b*-extension from an existing *p* ∈ *P*. If yes, then we extend *p*, so as to lengthen the chain that it forms with *p*_a_. If there are multiple collinear walks that reach *w*, we take the nearest one. If no, then we start a new collinear walk using just *x*. Figure 6 shows an example of updating a collinear walk and Figure 2 shows a full run of the algorithm for a single seed.

**Figure 6:**
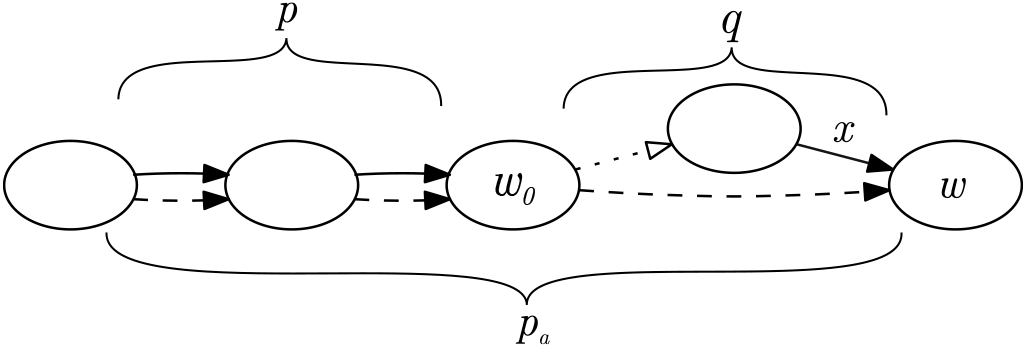
Illustration for Algorithm 2. A collinear walk *p* (solid) requires an update after the carrying path *p*_a_ is extended with the dashed edge (*w*_0_*, w*). The path *p*_a_ now ends at the vertex *w*, which has another incoming edge *x*. Since *x* is a part of *b*-extension of *p* (denoted by *q*), *p* can be appended with *q* to form a longer chain and boost the collinearity score.

Our description here only considers extending the initial seed to the right, i.e. using out-going edges in the graph. However, we also run the procedure to extend the initial seed to the left, using the in-coming edges. The case is symmetric and we therefore omit the details.

### 4.4 Other considerations

For simplicity of presentation, we have described the algorithm in terms of the ordinary de Bruijn graph; however, it is crucial for running time and memory usage that the graph is compacted first. Informally, the compacted de Bruijn graph replaces each non-branching path with a single edge. Formally, the vertex set of the compacted graph consist of vertices of the regular de Bruijn graph that have at least two outgoing (or ingoing) edges pointing at (incoming from) different vertices. Such vertices are called junctions. Let *ℓ* = *v*_1_*, …, v_n_* be the list of *k*-mers corresponding to junctions, in the order they appear in the underlying string *s*. The edge set of the compacted graph consists of edges {*v*_1_ → *v*_2_*, v*_2_ → *v*_3_*, …, v_n_*_−1_ → *v_n_*}. We efficiently construct the compacted graph using our previously published algorithm TwoPaCo (Minkin *et al.*, 2017).

This transformation maintains all the information while greatly reducing the number of edges and vertices in the graph. This makes the data structures smaller and allows the algorithm to fast-forward through non-branching paths, instead of considering each (*k* + 1)-mer one by one. Our previous description of the algorithm remains valid, except that the data structures operate with vertices and edges from the compacted graph instead of the ordinary one. The only necessary change is that when we look for an edge *y* parallel to *x*, we must also check that *y* and *x* spell the same sequence. This is always true in an ordinary graph but not necessarily in a compacted graph.

An important challenge of mammalian genomes is that they contain high-frequency (*k*+1)-mers, which can clog up our data structures. To handle this, we modify the algorithm by skipping over any junctions that correspond to *k*-mers occurring more than *a* times; we call *a* the abundance pruning parameter. Specifically, prior to constructing the edge set of the compacted de Bruijn graph, we remove all high abundance junctions from the vertex set. The edge set is constructed as before, but using this restricted list of junctions as the starting point. This strategy offers a way to handle high-frequency repeats at the expense of limiting our ability to detect homologous blocks that occur more than *a* times.

The organization of our data in memory is instrumental to achieving high performance. To represent the graph, we use a standard adjacency list representation, annotated with position information and other relevant data. We also maintain a list of the junctions in the paragraph above in the order they appear in the input sequences, thereby supporting next() queries. The walks in the collinear block *P* are stored as a dynamic sorted set, implemented as a binary search tree. The search key is the genome/position for the end of each walk. This allows performing binary search in line 2 of Algorithm 2.

Another aspect that we have ignored up until now is that DNA is double-stranded and collinear walks can be reverse-complements of each other. If *s* is a string, then let 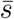 be its reverse complement. We handle double strandedness in the natural way by using the comprehensive de Bruijn graph, which is defined as 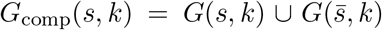 (Minkin *et al.*, 2017). Our algorithm and corresponding data structures can be modified to work with the comprehensive graph with a few minor changes which we omit here.

Our implementation is parallelized by exploring multiple seeds simultaneously, i.e. parallelizing the for loop at line 3 of Algorithm 1. This loop is not embarrassingly parallelizable, since two threads can start exploring two seeds belonging to the same carrying path. In such a case, there will be a collision on the data structure used to store used marks. To address this issue, we process the seeds in batches of fixed size. All the seeds within a batch are explored in parallel and the results are saved without modifying the “used” marks. Once the batch is processed, a single arbiter thread checks if there is any overlap in the used marks of the different threads. If there is, it identifies the sources of the conflict and reruns the algorithm at the conflicting seeds serially. Since most seeds do not yield valid carrying paths, such conflicts are rare. Once there is no conflict, the arbiter updates the used main data structures with the results of the batch. This design allows the computation result to be deterministic and independent of the number of threads used.

## Author contributions

Conceptualization, IM; Methodology, IM and PM; Software, IM; Validation, IM and PM, Writing - Original Draft, IM and PM; Writing - Review & Editing, IM and PM, Funding Acquisition, PM.

## Competing Interests

The authors declare no competing interests.

## Data availability

Table 4 contains the list of GenBank accession numbers of the mice genomes we used in our experiments (Figures 3, 4, Supplementary Figures S1, S2, S3). The nine simulated datasets we generated (Supplementary Figures S4, S5, S6), ground-truth alignments for the mouse data (Figure 4, Supplementary Figures S1, S2, S3), and alignments produced by SibeliaZ and Progressive Cactus (Figure 4, Supplementary Figures S2, S3) are available for download at https://github.com/medvedevgroup/SibeliaZ/blob/master/DATA.txt.

**Table 4:** Accession numbers of the assembled mice genomes available at GenBank.

| Strain | Accession number |
| --- | --- |
| C57BL/6J | GCA_000001635.8 [https://www.ncbi.nlm.nih.gov/assembly/GCA_000001635.8] |
| 129S1/SvImJ | GCA_001624185.1 [https://www.ncbi.nlm.nih.gov/assembly/GCA_001624185.1] |
| A/J | GCA_001624215.1 [https://www.ncbi.nlm.nih.gov/assembly/GCA_001624215.1] |
| AKR/J | GCA_001624295.1 [https://www.ncbi.nlm.nih.gov/assembly/GCA_001624295.1] |
| CAST/EiJ | GCA_001624445.1 [https://www.ncbi.nlm.nih.gov/assembly/GCA_001624445.1] |
| CBA/J | GCA_001624475.1 [https://www.ncbi.nlm.nih.gov/assembly/GCA_001624475.1] |
| DBA/2J | GCA_001624505.1 [https://www.ncbi.nlm.nih.gov/assembly/GCA_001624505.1] |
| FVB/NJ | GCA_001624535.1 [https://www.ncbi.nlm.nih.gov/assembly/GCA_001624535.1] |
| NOD/ShiLtJ | GCA_001624675.1 [https://www.ncbi.nlm.nih.gov/assembly/GCA_001624675.1] |
| NZO/HiLtJ | GCA_001624745.1 [https://www.ncbi.nlm.nih.gov/assembly/GCA_001624745.1] |
| PWK/PhJ | GCA_001624775.1 [https://www.ncbi.nlm.nih.gov/assembly/GCA_001624775.1] |
| WSB/EiJ | GCA_001624835.1 [https://www.ncbi.nlm.nih.gov/assembly/GCA_001624835.1] |
| BALB/cJ | GCA_001632525.1 [https://www.ncbi.nlm.nih.gov/assembly/GCA_001632525.1] |
| C57BL/6NJ | GCA_001632555.1 [https://www.ncbi.nlm.nih.gov/assembly/GCA_001632555.1] |
| C3H/HeJ | GCA_001632575.1 [https://www.ncbi.nlm.nih.gov/assembly/GCA_001632575.1] |
| LP/J | GCA_001632615.1 [https://www.ncbi.nlm.nih.gov/assembly/GCA_001632615.1] |

## Code availability

Our tool is open source and freely available at https://github.com/medvedevgroup/SibeliaZ.

## Acknowledgements

We would like to thank Mikhail Kolmogorov for useful suggestions on the empirical evaluation of our algorithm; Robert Harris for his help with running MultiZ; Son Pham for introducing us to the problem; and the Ensembl support team for helping us with retrieving the gene annotations.

This work has been supported in part by NSF awards DBI-1356529, CCF-1439057, IIS-1453527, and IIS-1421908 to PM. Research reported in this publication was supported by the National Institute Of General Medical Sciences of the National Institutes of Health under Award Number R01GM130691. The content is solely the responsibility of the authors and does not necessarily represent the official views of the National Institutes of Health.

## Supplementary Notes, Figures, and Tables

### Supplementary Note 1 Other related work

A closely related problem to multiple whole-genome alignment is synteny reconstruction. In this setting, genomes are decomposed into large blocks such that the gene order within each block is preserved. This is similar to locally collinear blocks, but collinear blocks are usually smaller blocks representing single genes or exons (or non-coding DNA). Collinear blocks can be viewed as high resolution synteny blocks and, in general, the distinction between the two concepts can be blurry. For a discussion on representation of synteny blocks at multiple scales, see Minkin *et al.* (2013b). Synteny blocks are often reconstructed from anchors such as genes (Pevzner and Tesler, 2003a; Ng *et al.*, 2009; Pham and Pevzner, 2010; Proost *et al.*, 2011) and, less commonly, from the nucleotide sequences directly (Minkin *et al.*, 2013b; Doerr and Moret, 2018).

A related active research area is data structures for representing pan-genomes (Tettelin *et al.*, 2005). A pan-genome as a collection of related genomes that are to be analyzed jointly. For a review on computational pan-genomics, see (Vernikos *et al.*, 2015; Marschall *et al.*, 2018; Zekic *et al.*, 2018). Constructing a data structure for the efficient storage and querying of a pan-genome is related but tangential to the problem of identifying collinear blocks, which we consider in this paper. Pan-genome data structures are concerned with efficiently representing the homology within the pan-genome, while we focus on fast algorithms for obtaining such homologies. There is naturally some overlap between the two areas, e.g. some pan-genome tools include a multiple whole-genome alignment component (Ernst and Rahmann, 2013; Laing *et al.*, 2010). Others use the de Bruijn graph for representing the pan-genome (Marcus *et al.*, 2014; Holley *et al.*, 2016; Beller and Ohlebusch, 2016; Sheikhizadeh *et al.*, 2016). Our approach also relies on a de Bruijn graph, though we use it as a technique to find collinear blocks rather than to represent them.

### Supplementary Note 2 Parameter details and command lines

We tried to find the optimal parameters for all tools. For Sibelia, which could only run on simulated data, we used the parameter set designed to yield the highest sensitivity (called the “far” set in Sibelia). Progressive Cactus requires a phylogenetic tree in addition to the input genome which it uses for adjusting the internal parameters. For the simulated datasets, we used the real tree generated by the simulator; for the mice genomes, we used the guide tree from (Lilue *et al.*, 2018). Multiz+TBA were run with default parameters since its documentation does not provide a clear guideline on how to adjust the parameters according to evolutionary distances between the input sequences. We could not compile the version of MultiZ+TBA publicly available for download and used a slightly modified version provided by Robert S. Harris. For TwoPaCo and spoa, we set the parameters following the guidelines provided with the respective software. SibeliaZ was run with *k* = 25, *b* = 200, *m* = 50, and *a* = 150.

We performed all experiments on a machine running Ubuntu 16.04.3 LTS with 512 GB of RAM and a 64 core CPU Intel Xeon CPU E5-2683 v4. We were limited to using at most 32 threads at any given time. Progressive Cactus was run with 32 threads, since the authors recommended to use as many threads as possible for the best performance. MultiZ+TBA and Sibelia are both single-threaded. (There were several submissions to Alignathon which used an extensively parallelized MultiZ or TBA; unfortunately, the software packages used for those submissions are not available publicly for download.) TwoPaCo and SibeliaZ-LCB were run 16 and 32 threads respectively. We note that spoa is run on each block, and our software includes a wrapper to automate this.

Here are the exact command lines for the tools we ran.

TwoPaCo:

~~~
twopaco -k <k_value> -f <bloom_filter_size> -t 16 -o <dbg_graph> <genomes_file>
SibeliaZ-LCB:
SibeliaZ-LCB --fasta <genomes_file> --graph <dbg_graph> -o <output_directory>
-k 25 -b 200 -m 50 -a 150 -t 4
spoa:
spoa <input_fasta_file> -l 1 -r 1 -e 8
Sibelia:
Sibelia <genomes_file> -o <output_directory> -s far --lastk 50 -m 50 --nopostprocess
MultiZ:
all_bz <guide_tree>
tba <guide_tree> *.*.maf <outputMafFile>
~~~

Progressive Cactus:

~~~
runProgressiveCactus.sh --maxThreads 32 <seqFile> <workDir> <outputHalFile> source ./environment && hal2mafMP.py <outputHalFile> <outputMafFile>
~~~

All running times and memory usage numbers were obtained using the GNU time utility. The exact versions of the software are in Table S4.

### Supplementary Note 3 Simulation details

We used small simulated data in order to understand the role of a dataset’s genomic distance and of our parameter settings. We used ALF (Dalquen *et al.*, 2011) for simulation because it simulates point mutations as well as genome-wide events such as inversions, translocations, fusions/fissions, gene gain/loss, and lateral gene transfer. Furthermore, ALF is useful for benchmarking as it also produces an alignment which represents the true homology between the genomes, making it possible to directly assess the precision and recall. We simulated 6 datasets, each one consisting of 10 bacterial genomes. Each genome is composed of 1500 genes and of size approximately 1.5 Mbp. We used such relatively small datasets to allow us to efficiently explore the parameter space. Each of the 6 datasets corresponded to a different parameter for distance from the root to leaf species, which we varied from 0.03 to 0.18 substitutions per site with the step of 0.03. In ALF, different proteins evolve with different rates, which are derived from the base value using a probabilistic distribution. See *Dalquen et al.* (2011) for more details and the simulation recipes for the exact values of the parameters. For genome-wide events, we used ALF’s default rates. Links to download the the simulation parameter files, the simulated genomes, and their alignments are available at the GitHub repository (see Section 4.4 section in the main paper).

**Supplementary Figure S1:**
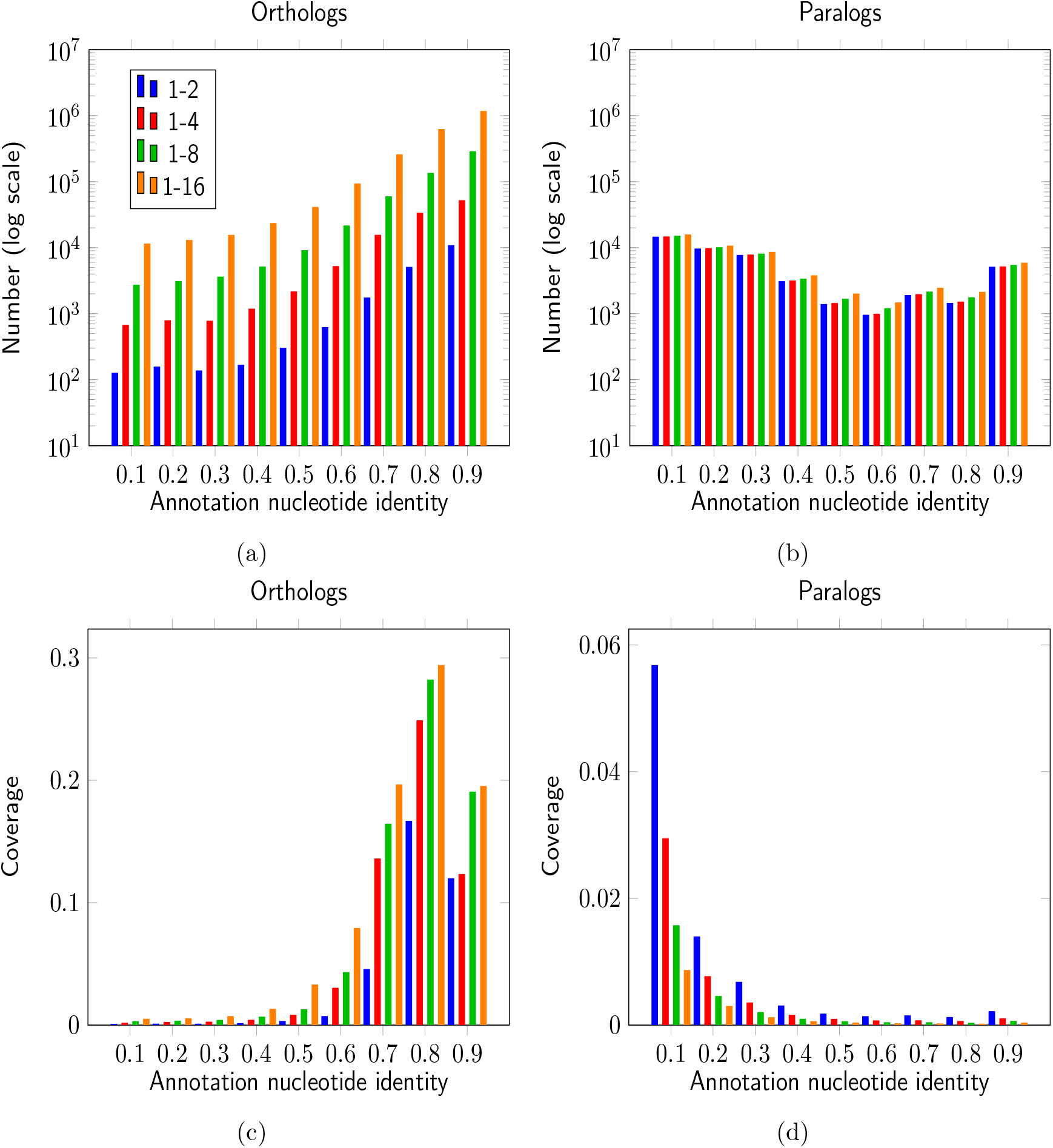
Properties of the pairwise alignments constructed from pairs of homologous protein-coding genes in the various mice datasets. Panels (a) and (b) show the total number of genes for each nucleotide identity value for orthologous and paralogous pairs respectively; (c) and (d) demonstrate coverage of the genomes by these two categories of the genes.

**Supplementary Figure S2:**
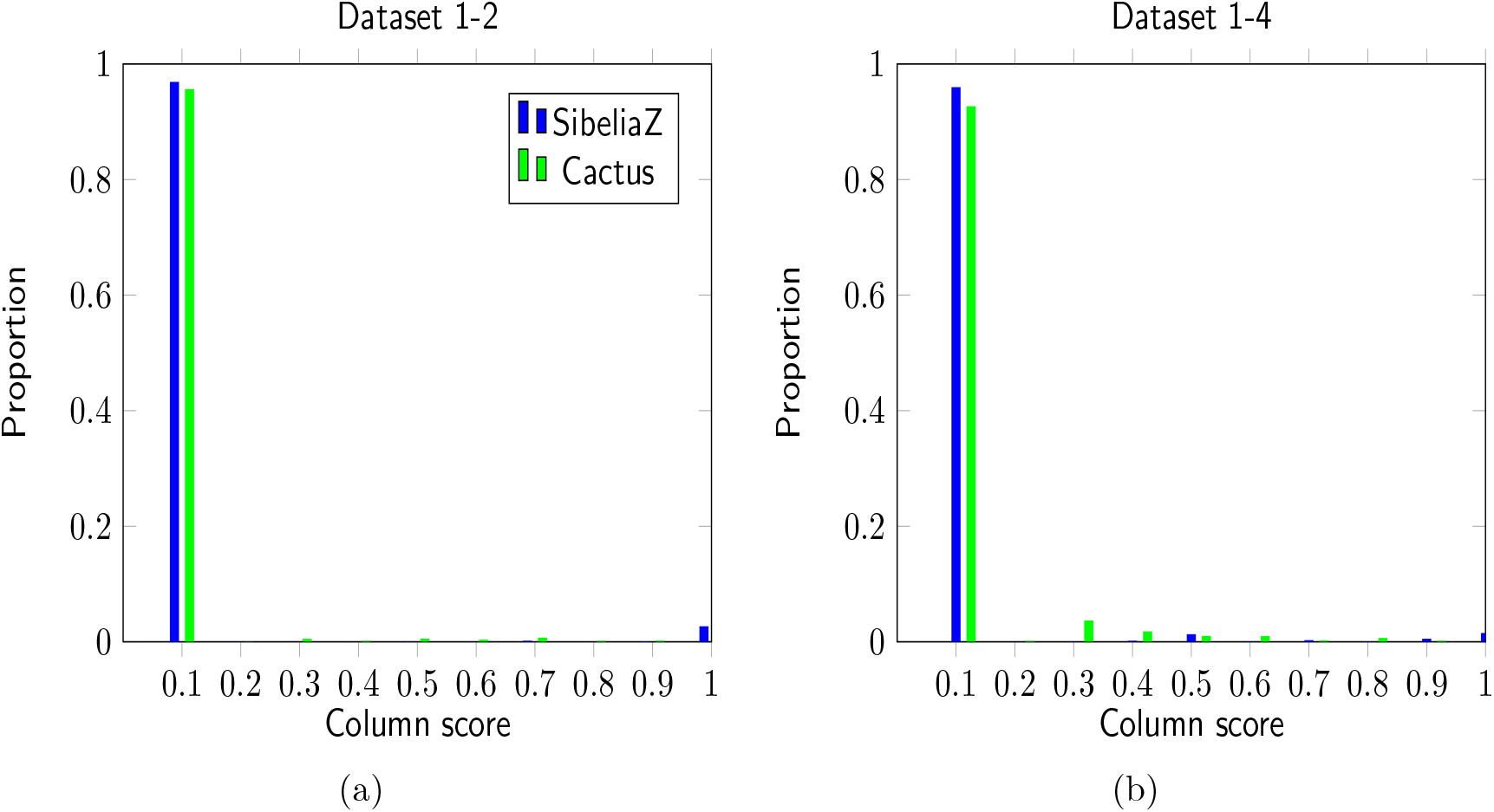
Histogram of the average number of nucleotide differences *π*(*c*) calculated for each column of SibeliaZ’s and Cactus’ alignment, for datasets 1-2 (a) and 1-4 (b) consisting of 2 and 4 mice geneomes respectively.

**Supplementary Figure S3:**
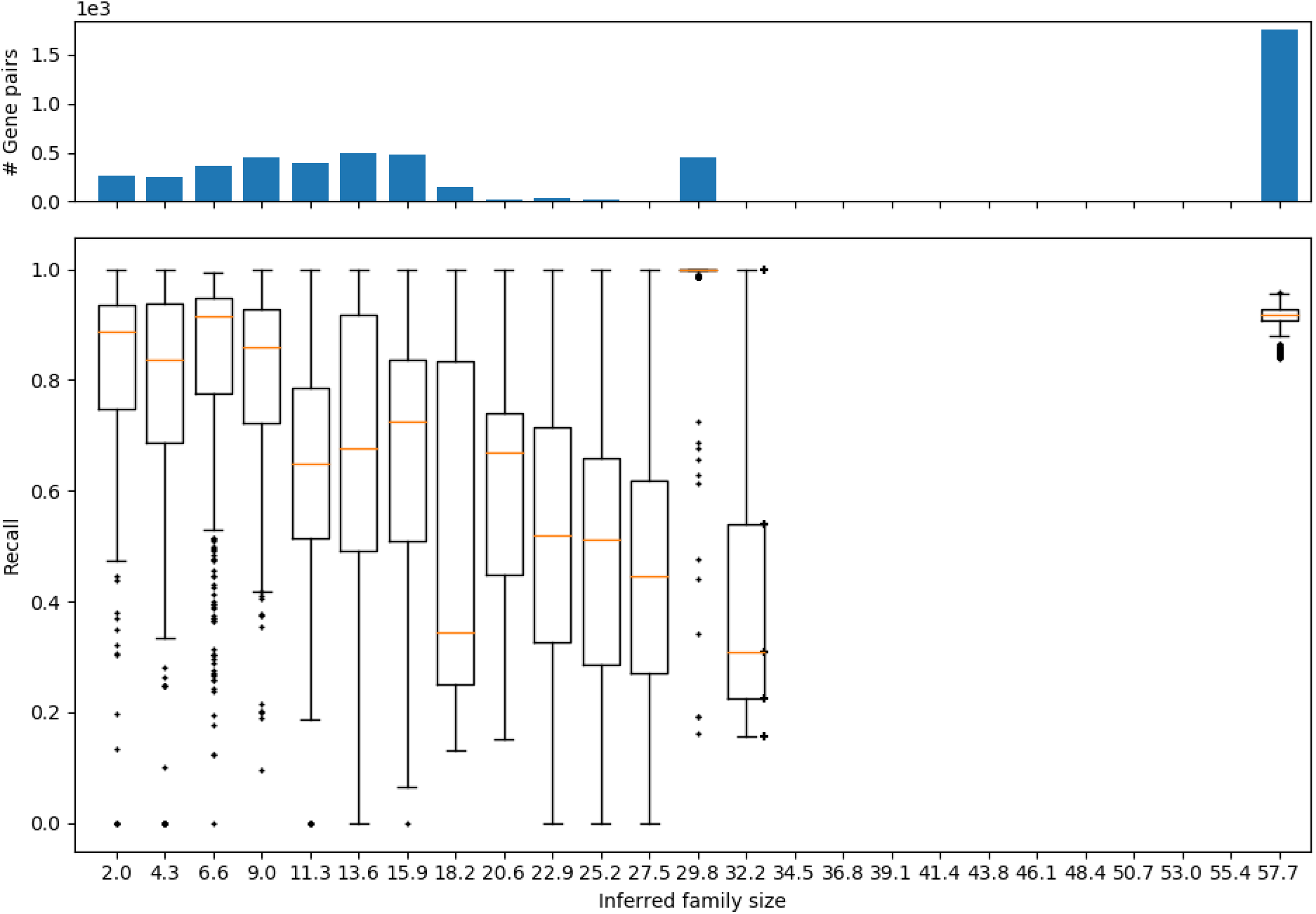
The recall of paralogous genes by SibeliaZ as a function of inferred family size, using the two-mice dataset. The *n* = 5152 gene pairs were split into 25 equally-sized disjoint bins based on the inferred family size. The top histogram shows the number of gene pairs in each bin. Exact sizes for each bin are (267, 250, 360, 452, 396, 489, 481, 155, 24, 35, 21, 12, 450, 5, 0, 0, 0, 0, 0, 0, 0, 0, 0, 0, 1755). Points belonging to non-empty bins of size less than 10 are shown individually. Each box plot shows the median (middle line), the interquartile range (outer borders of the box), minimum and maximum values within ±1.5 of the interquartile range (whiskers). Data points outside of the ±1.5 interquartile range are represented by individual data points.

**Supplementary Figure S4:**
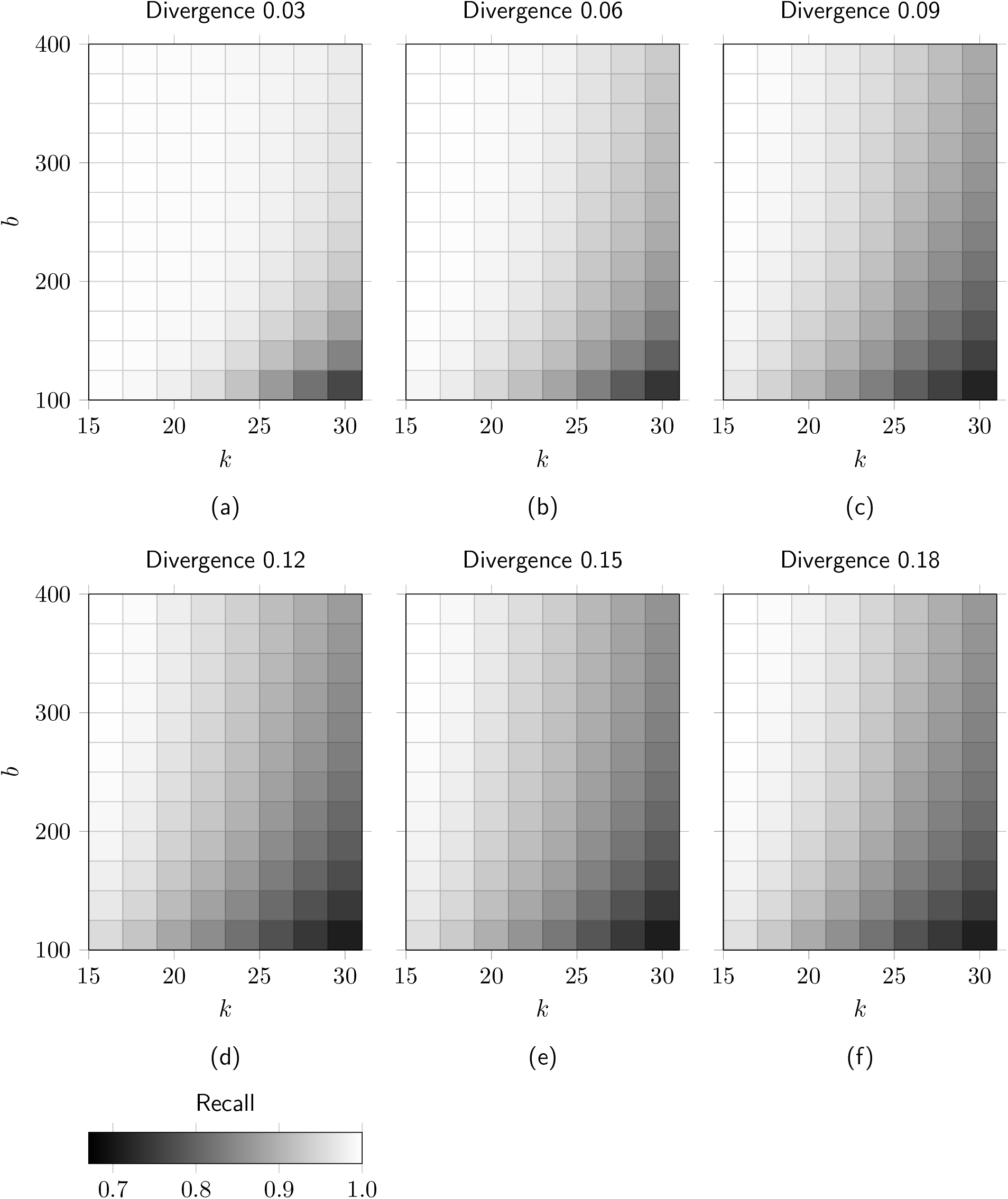
Effects of the parameters *k* and *b* on recall. Each panel (a-f) contains a heatmap corresponding to a simulated dataset with the specified root-to-leaf divergence in substitutions per site and a cell corresponds to a combination of parameters.

**Supplementary Figure S5:**
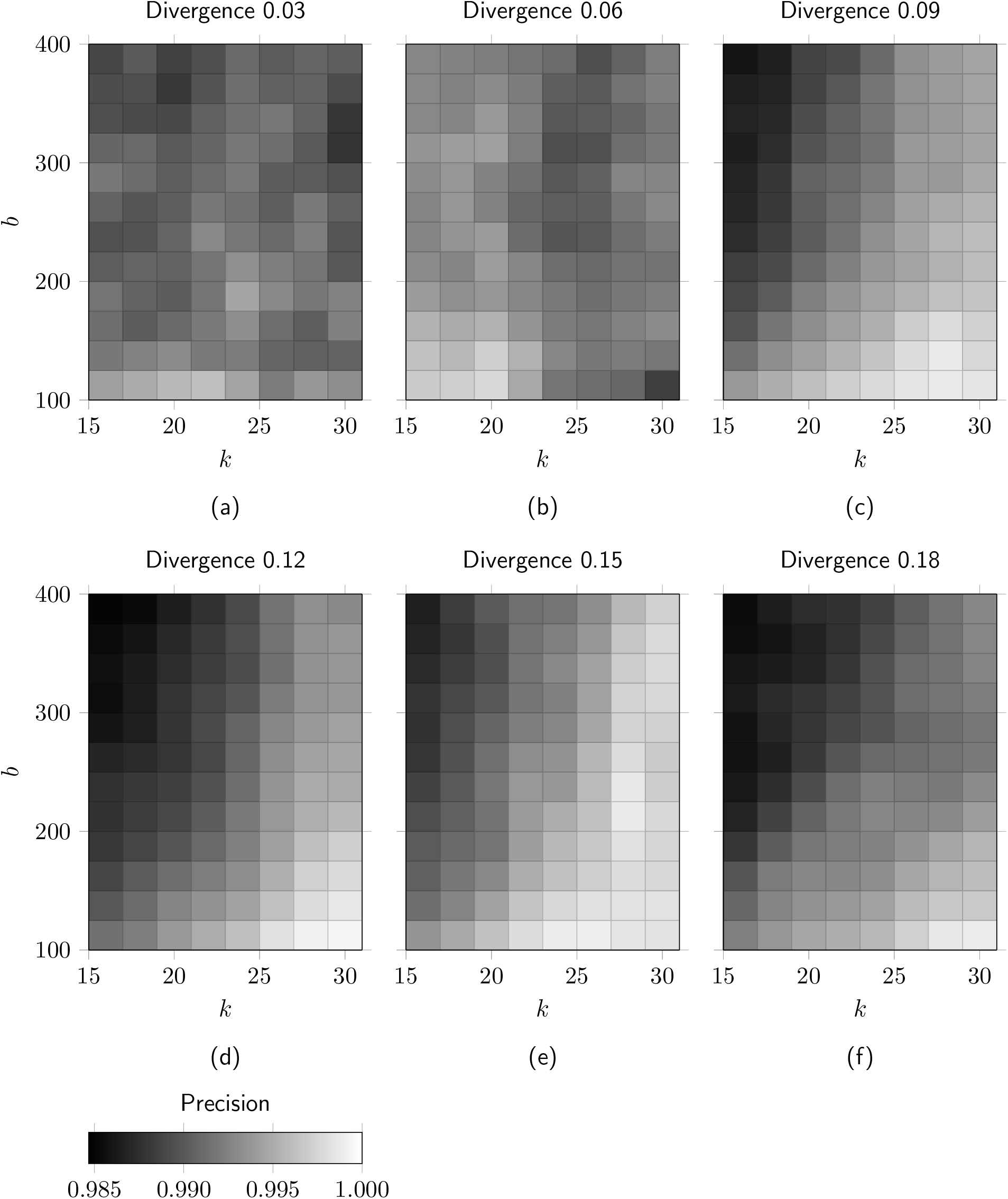
Effects of the parameters *k* and *b* on precision. Each panel (a-f) contains a heatmap corresponding to a simulated dataset with the specified root-to-leaf divergence in substitutions per site and a cell corresponds to a combination of parameters.

**Supplementary Figure S6:**
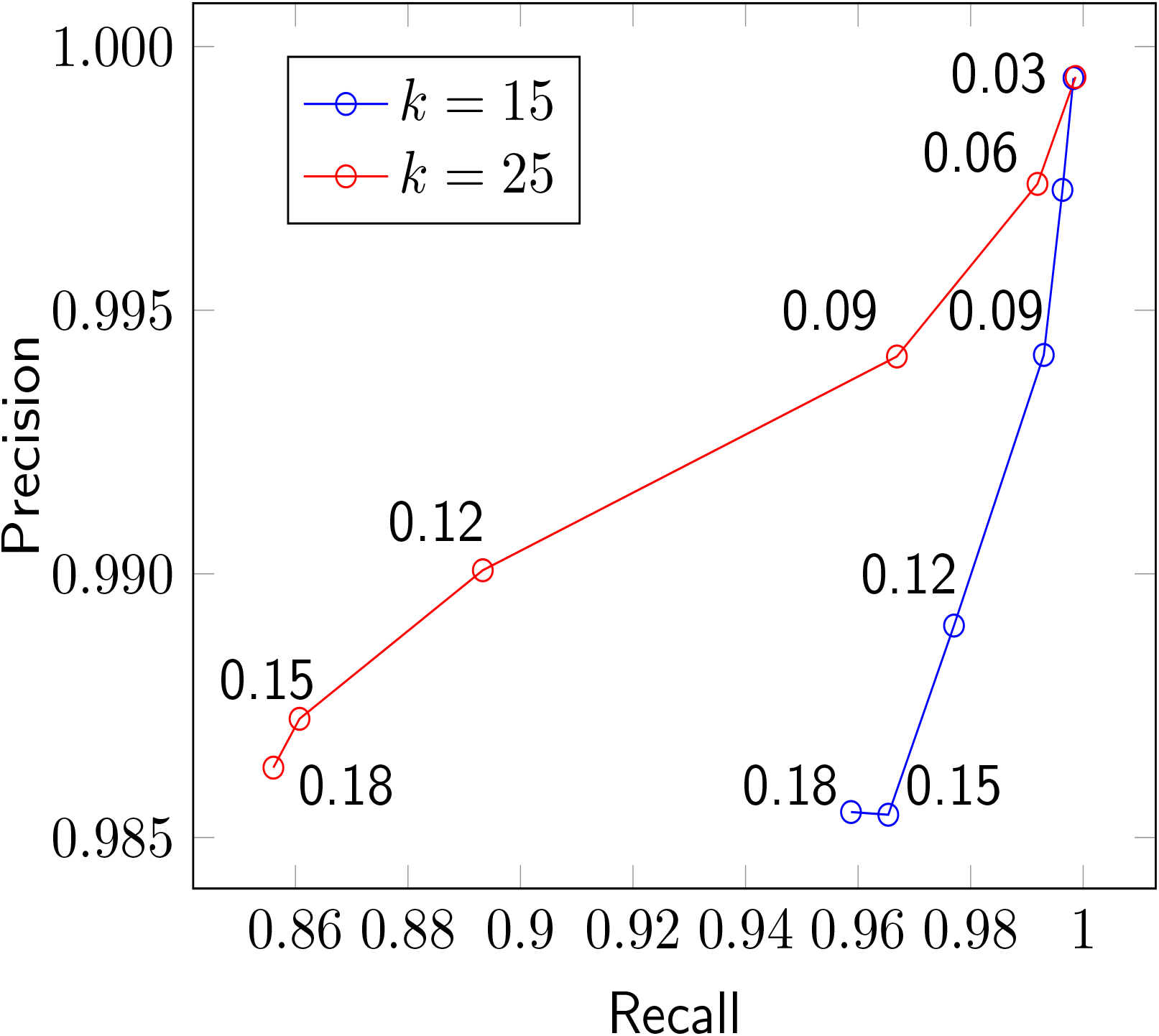
The accuracy of SibeliaZ as a function of genomic divergence. Each point is labeled with the height of the phylogenetic tree (in terms of substitutions per site) of its respective simulated dataset.

**Supplementary Table S1:** Properties of the assembled mice genomes available at GenBank.

| ID | Strain | Size (Mb) | N. Scaffolds |
| --- | --- | --- | --- |
| 1 | C57BL/6J | 2,819 | 336 |
| 2 | 129S1/SvImJ | 2,733 | 7,154 |
| 3 | A/J | 2,630 | 4,688 |
| 4 | AKR/J | 2,713 | 5,953 |
| 5 | CAST/EiJ | 2,654 | 2,977 |
| 6 | CBA/J | 2,922 | 5,466 |
| 7 | DBA/2J | 2,606 | 4,105 |
| 8 | FVB/NJ | 2,589 | 5,013 |
| 9 | NOD/ShiLtJ | 2,982 | 5,544 |
| 10 | NZO/HiLtJ | 2,699 | 7,022 |
| 11 | PWK/PhJ | 2,560 | 3,140 |
| 12 | WSB/EiJ | 2,690 | 2,239 |
| 13 | BALB/cJ | 2,627 | 3,825 |
| 14 | C57BL/6NJ | 2,807 | 3,894 |
| 15 | C3H/HeJ | 2,701 | 4,069 |
| 16 | LP/J | 2,731 | 3,499 |

**Supplementary Table S2:** Running time (minutes) and memory usage (gigabytes, in parenthesis) of SibeliaZ and Cactus on the mice datasets. A dash in a column indicates that the program did not complete within in a week.

| Dataset | SibeliaZ/TwoPaCo | SibeliaZ/SibeliaZ-LCB | SibeliaZ/spoa | SibeliaZ/Total | Cactus |
| --- | --- | --- | --- | --- | --- |
| 1-2 | 12 (9.30) | 74 (36.00) | 68 (121.50) | 154 (121.50) | 2,279 (37.50) |
| 1-4 | 25 (17.70) | 96 (72.70) | 115 (133.60) | 236 (133.60) | 6,105 (89.80) |
| 1-8 | 49 (34.50) | 104 (106.30) | 240 (132.50) | 393 (132.50) | - |
| 1-16 | 101 (68.10) | 153 (183.60) | 736 (133.50) | 990 (183.60) | - |

**Supplementary Table S3:** Recall and precision of SibeliaZ and Cactus. We used pairwise local alignments of chromosomes 1 from the different datasets produced by LASTZ as the ground truth.

| Dataset | SibeliaZ Recall | Cactus Recall | SibeliaZ Precision | Cactus Precision |
| --- | --- | --- | --- | --- |
| 1-2 | 0.95 | 0.97 | 0.93 | 0.89 |
| 1-4 | 0.95 | 0.92 | 0.96 | 0.90 |

**Supplementary Table S4:**
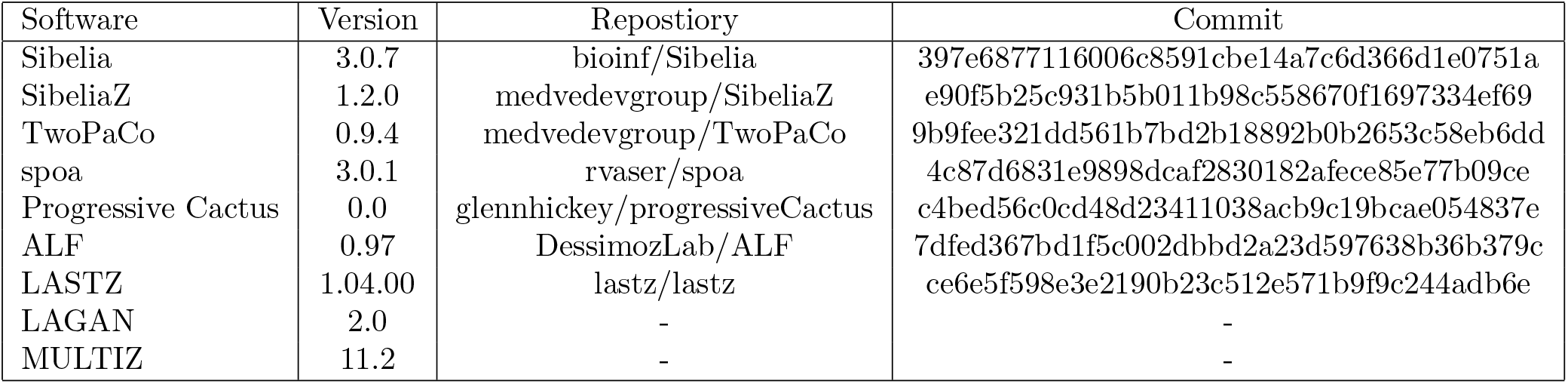
GitHub revisions of the software we used. We used the most up-to-date versions available at the time of development of our project. A dash indicates that we downloaded the software from the author’s website.

## Notes

https://github.com/medvedevgroup/SibeliaZ

### Competing Interest Statement

The authors have declared no competing interest.

### Summary of Updates

We have updated the design of our experiments to make benchmarking fair. In addition, we improved the presentation of our method.

